# The immunomodulatory dCache chemoreceptor TlpA of *Helicobacter pylori* binds multiple attractant and antagonistic ligands via distinct sites

**DOI:** 10.1101/2021.04.07.438267

**Authors:** Kevin S. Johnson, Bassam Elgamoudi, Freda E.-C. Jen, Christopher Day, Emily Goers Sweeney, Megan L. Pryce, Karen Guillemin, Thomas Haselhorst, Victoria Korolik, Karen M. Ottemann

## Abstract

The *Helicobacter pylori* chemoreceptor TlpA plays a role in dampening host inflammation during chronic stomach colonization. TlpA has a periplasmic dCache_1 domain, a structure that is capable of sensing many ligands; however, the only characterized TlpA signals are arginine, bicarbonate, and acid. To increase our understanding of TlpA’s sensing profile, we screened for diverse TlpA ligands using ligand binding arrays. TlpA bound seven ligands with affinities in the low to middle micromolar ranges. Three of these ligands, arginine, fumarate, and cysteine, were TlpA-dependent chemoattractants, while the others elicited no response. Molecular docking experiments, site-directed point mutants, and competition surface plasmon resonance binding assays suggested that TlpA binds ligands via both the membrane-distal and -proximal dCache_1 binding pockets. Surprisingly, one of the non-active ligands, glucosamine, acted as a chemotaxis antagonist, preventing the chemotaxis response to chemoattractant ligands and acted to block binding of ligands irrespective of whether they bound the membrane-distal or -proximal dCache_1 subdomains. In total, these results suggest TlpA senses multiple attractant ligands as well as antagonist ones, an emerging theme in chemotaxis systems.

**Importance:** Numerous chemotactic bacterial pathogens depend on the ability to sense a diverse array of signals through chemoreceptors to achieve successful colonization and virulence within their host. The signals sensed by chemoreceptors, however, are not always fully understood. This is the case for TlpA, a dCache_1 chemoreceptor of *H. pylori* that enables the bacteria to induce less inflammation during chronic infections. *H. pylori* causes a significant global disease burden, which is driven by the development of gastric inflammation. Accordingly, it is essential to understand the processes by which *H. pylori* modulates host inflammation. This work uncovers the signals that TlpA can sense and highlights the underappreciated ability for regulating chemotactic responses by antagonistic chemoreceptor ligands, which is an emerging theme among other chemotactic systems.

## Introduction

Chemotaxis is a vital host colonization strategy used by many pathogens, including *Helicobacter pylori, Campylobacter jejuni, Borrelia burgdorferi, Pseudomonas aeruginosa, Vibrio cholerae*, and *Salmonella enterica*. How chemotaxis benefits bacteria, however, varies. Pathogens have been found to use chemotaxis to access growth-promoting nutrients, locate signaling molecules that regulate virulence gene expression, spread throughout tissue and into specific niches, and affect host interactions that control inflammation (Matilla and Krell, 2018). Chemotaxis signaling systems are highly conserved, and their widespread presence in pathogens underscores the importance of understanding their roles in colonization (Matilla and Krell, 2018; Wuichet and Zhulin, 2010).

One pathogen that requires chemotaxis for multiple infection aspects is *H. pylori*. This Gram-negative bacterium chronically colonizes the stomach of nearly half of the world’s population and ∼35% of individuals in the United States (Hooi et al., 2017). Stomach colonization results in chronic inflammation, and a subset of individuals develop ulcers and gastric cancer (Cover and Blaser, 2009; Plummer et al., 2015). *H. pylori* presents a significant disease burden, with ∼700,000 deaths from gastric cancer yearly (Ferlay et al., 2015). While many people are infected, the degree of host inflammation varies, which ultimately drives disease severity (Arnold et al., 2011; White et al., 2015; Wroblewski et al., 2010). We understand some *H. pylori* properties that dictate inflammation severity, such as the Cag pathogenicity island (Blosse et al., 2018; Javed et al., 2019). Still, the full compendium of *H. pylori* properties that modulate this host response is not yet understood.

*H. pylori* chemotaxis has been linked to host inflammation (Johnson and Ottemann, 2018; McGee et al., 2005; Rolig et al., 2011; Williams et al., 2007). Specifically, mutants missing key chemotaxis signal transduction proteins trigger less host inflammation, despite achieving normal colonization levels, while mutants missing either the chemoreceptors TlpA or TlpB cause elevated inflammation (McGee et al., 2005; Rolig et al., 2011; Williams et al., 2007). Chemoreceptors head the chemotaxis signal transduction system and dictate which signals a bacterium responds to. Loss of individual chemoreceptors within a system alters a bacterium’s sensing profile but does not cause a complete loss of chemotactic ability, presumably biasing the bacterium towards signals sensed by the remaining chemoreceptors.

*H. pylori* possesses four chemoreceptors—TlpA, TlpB, TlpC, and TlpD. Each of these plays non-identical roles in infection. TlpA, C, and D are required for colonization, while TlpA and B are required for inflammation control (Andermann et al., 2002; Rolig et al., 2012; Williams et al., 2007). In this work, we focus on TlpA, which plays multiple roles promoting early but not late colonization and dampening later inflammation. At early times, 2 weeks post-infection, *H. pylori ΔtlpA* displays a modest colonization defect compared to wild type (WT) as the sole infecting strain, a deficiency exacerbated by WT coinfection (Andermann et al., 2002; Rolig et al., 2012; Williams et al., 2007). However, during the chronic stage of infection after 6 months, *H. pylori ΔtlpA* colonize to normal levels but induce significantly more histologically evident inflammation than WT (Williams et al., 2007).

TlpA is a transmembrane chemoreceptor with a periplasmic double Cache (dCache_1) ligand-binding domain (LBD) (Sweeney et al., 2018; Upadhyay et al., 2016). Cache domains are ubiquitous extracellular sensing domains found in both eukaryotes and prokaryotes, where they are the most common extracellular sensing domain (Upadhyay et al., 2016). Cache domains bind a wide variety of small molecules, but mostly amino acids, modified amino acids, and carboxylic acids (Upadhyay et al., 2016). Many Cache domains have been found to bind multiple ligands (Upadhyay et al., 2016). dCache_1 domains have two Cache subdomains, a membrane-distal and -proximal subdomain, which each can bind ligands, although most commonly the ligands are bound in the membrane-distal domain (Machuca et al., 2017; Mckellar et al., 2015; Upadhyay et al., 2016). TlpA has some identified chemotaxis-active ligands, including arginine and sodium bicarbonate (Cerda et al., 2003, 2011). Additionally, TlpA has been shown to play a subtle role in sensing acidic pH, but to a much lesser extent than TlpB or TlpD (Huang et al., 2017). Whether TlpA senses additional ligands or how any of these ligands are bound, however, is unknown.

Given the sensing potential of dCache_1 chemoreceptors, we hypothesized that TlpA would be capable of sensing ligands beyond those previously reported. Knowing a full set of ligands is critical for interpreting the TlpA-associated phenotypes. In this study, we identified new TlpA ligands, and characterized their binding and ability to induce a chemotactic response. Ligand binding arrays were used to screen a broad set of ligands for binding to TlpA, resulting in the identification and verification of seven TlpA specific ligands. Use of a temporal chemotaxis assay enabled us to determine that three ligands, arginine, fumarate, and cysteine, acted as TlpA-sensed chemoattractants, while the other ligands elicited no response. Molecular modeling experiments, assessment of TlpA point mutants, and SPR competition assays suggested that TlpA ligands interact with two distinct sites. Furthermore, one of the high-affinity non-chemotactic TlpA ligands, glucosamine, blocked attractant responses to, and binding of, chemoattractant ligands, thus acting as an antagonist. Overall, our findings suggest TlpA responds to several key *H. pylori* nutrients using both dCache_1 subdomains, with some acting as agonists and some as antagonists for a chemotaxis attractant response.

## Results

### TlpA interacts directly with multiple ligands

TlpA’s LBD (TlpA_LBD_) interactions with potential ligands were assessed using small-molecule arrays containing amino acids, organic acids, salts, and glycans (Supplemental Table 1). TlpA_LBD_ bound seven small molecules: arginine, cysteine, fumarate, glucosamine, malic acid, thiamine, and alpha-ketoglutarate (Table 1 and Supplemental Figure 1). Glycan arrays, containing both simple and complex glycans were also interrogated but no binding was detected.

**Table 1.** TlpA_LBD_ ligand binding analysis from ligand binding array screen and surface plasmon resonance. Data represents the mean values (± SD) of three independent experiments (n= 3). For ligand binding array, results reported as +, positive binding; +/-, intermediate binding; -, no binding. Binding affinity (µM) was determined by SPR.

| Ligand | Array | Binding affinity (μM) |
| --- | --- | --- |
| Arginine | + | $2 \pm 0.11$ |
| Cysteine | +/- | $4.7 \pm 0.3$ |
| Fumarate | + | $10 \pm 1.53$ |
| Glucosamine | + | $10.5 \pm 2.8$ |
| Malic Acid | +/- | 46 ± 17 |
| Thiamine | +/- | 60 ± 0.6 |
| Alpha-Ketoglutarate | + | 224 ± 11.2 |

We next determined TlpA_LBD_ ligand binding affinity by surface plasmon resonance (SPR). Arginine, cysteine, fumarate, and glucosamine all exhibited high-affinity binding (K_d_ <10 µM), while malic acid, thiamine, and alpha-ketoglutarate showed lower affinities (>45 µM) (Table 1). Overall, these data suggest that the TlpA_LBD_ can interact directly with these seven ligands, with affinities ranging from 2-224 µM.

### Some TlpA ligands act as chemoattractants while others elicit no response

*H. pylori* chemotactic responses towards the seven putative TlpA dependent ligands were examined by a live-cell video microscopy assay that measures the temporal chemotaxis response to test ligands. In this assay, attractants eliciting fewer direction changes, and repellents more direction changes compared to basal levels (Collins et al., 2016; Goers Sweeney et al., 2012; Lertsethtakarn et al., 2015; Machuca et al., 2017; Rader et al., 2011; Schweinitzer et al., 2008; Terry et al., 2006). Several TlpA ligands were acidic in solution, leading to the appearance of significant TlpA-independent chemorepellent responses; these were cysteine, thiamine, malic acid, and alpha-ketoglutarate (Supplemental Fig. 2A-C). Acidic conditions are sensed by chemoreceptors other than TlpA (Croxen et al., 2006; Goers Sweeney et al., 2012; Huang et al., 2017) and potentially mask chemotactic responses to the ligands being tested. Accordingly, the pH of the ligand stocks for cysteine, thiamine, malic acid, and alpha-ketoglutarate was neutralized using NaOH to match the pH of the water used in the mock-treated control. This treatment eliminated the confounding effect of media acidification on assessing chemotactic responses (Supplemental Fig. 2C). After incorporating these adjustments, we found that the addition of arginine, fumarate, or cysteine resulted in fewer direction changes for WT *H. pylori* (Fig. 1A, B, C), suggesting these compounds were attractants. The highest concentration tested, 10 mM, induced the most significant and robust attractant responses for each ligand (arginine, P<0.01; fumarate, P<0.01; cysteine, P<0.001) (Fig. 1A, B, C). Responses to 1 and 0.1 mM ligands were apparent but not significant when compared to the untreated control. Glucosamine, thiamine, malic acid, and alpha-ketoglutarate induced no significant direction changes at any concentration tested (Fig. 1D and E), suggesting they do not act as attractants or repellents.

**Figure 1.**
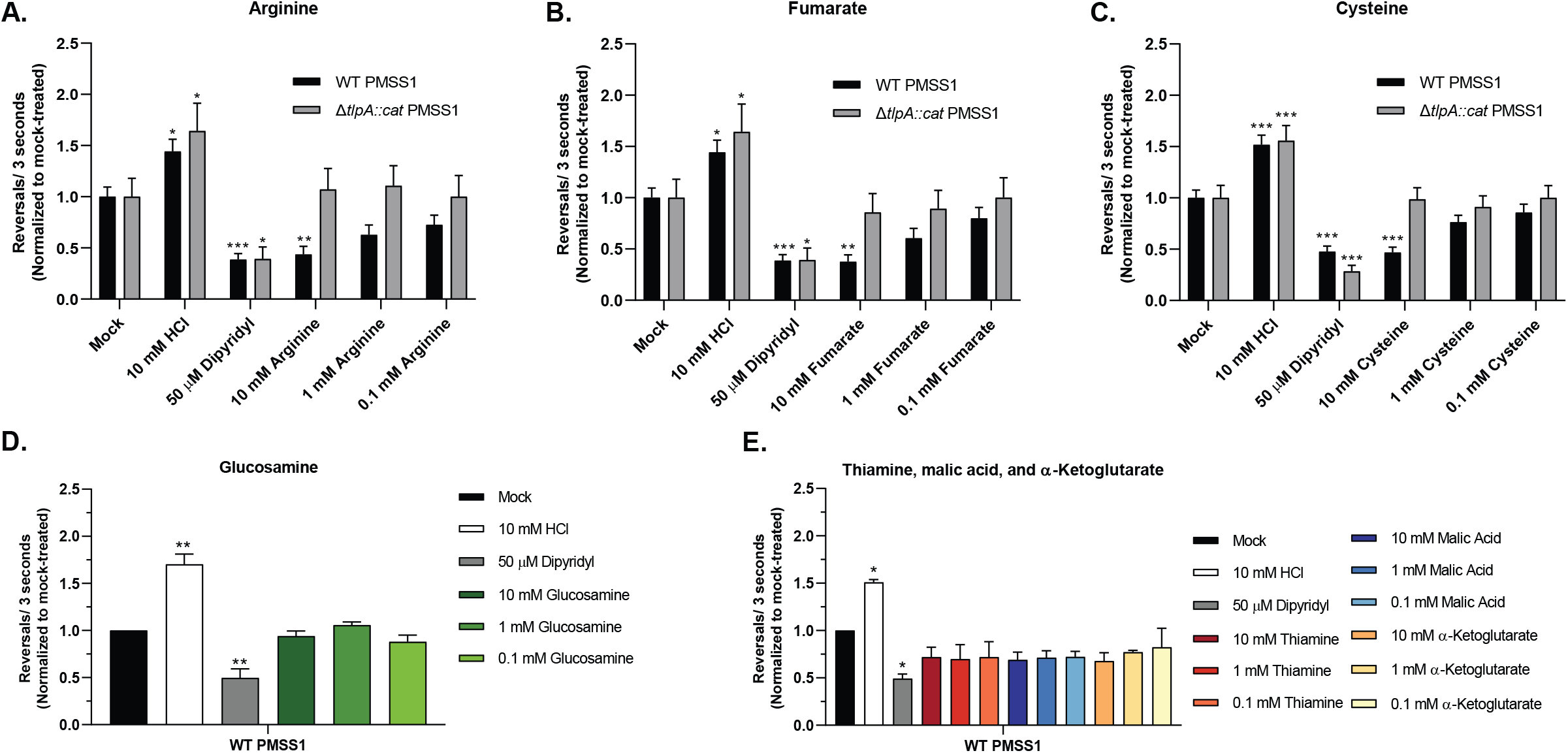
*H. pylori* responds to arginine, fumarate, and cysteine as TlpA-dependent chemoattractants in the temporal chemotaxis assay. Cultures of *H. pylori* PMSS1 WT and *ΔtlpA* were grown overnight, back-diluted then incubated until an OD_600_ of 0.12– 0.15 was reached. Cultures were treated with water (Mock) or with various concentrations of compounds as indicated. For Fig. 1C and E, the Ph of cysteine, thiamine, malic acid, and alpha-ketoglutarate stocks was adjusted using NaOH to match the pH of the water used for the untreated control. The cells were immediately filmed, and direction changes were counted over a 3 second swimming period in at least 100 cells per treatment from 3 biological replicates. Data is normalized to the untreated control for each strain, as described in the methods. Error bars represent the standard error of the mean. *, P<0.05; **, P<0.01; ***, P<0.001, comparisons to untreated control per strain using two-way ANOVA, Dunnett’s multiple comparison test.

To determine if chemoattractant responses toward arginine, fumarate, and cysteine were TlpA dependent, the same tracking experiments were repeated with a mutant lacking *tlpA* (Δ*tlpA*). The Δ*tlpA* mutant retained general chemotactic ability, producing a significant attractant and repellent response to controls dipyridyl and HCl, respectively, but failed to exhibit a chemotactic response to arginine, fumarate, or cysteine (Fig. 1A, B, C). These results suggest that the high-affinity ligands arginine, fumarate, and cysteine are TlpA-dependent chemoattractants.

### TlpA has at least two ligand interaction sites

To gain insight into how TlpA_LBD_ binds ligands, a blind docking modeling experiment was carried out using AutoDock Vina (Krieger et al., 2002; Wang et al., 2016). We focused on two high-affinity ligands, arginine and fumarate. We found that these two ligands occupied two main sites, referred to as clusters (Supplemental Table 2), which map to the membrane-distal and -proximal dCache_1 binding pockets (Fig. 2A and D). Arginine was placed mostly in Cluster D (55%, Supplemental Table 3), which is located in the membrane-distal dCache_1 domain. Arg-153 dominated this binding interaction with stabilization from Tyr-151 (Fig. 2B and C). Fumarate, in contrast, was placed mostly in Cluster A (45%, Supplemental Table 3), in the membrane-proximal dCache_1 domain, with Phe-203 being the most crucial residue required for interaction with fumarate (Fig. 2E and F). The docking experiment further revealed that although fumarate and arginine are likely to have two distinct preferred binding sites, they both can bind to their reciprocal sites. For example, 25% of the models had arginine found in fumarate’s preferred Cluster A (Supplemental Table 3). Overall, these analyses suggest that arginine is more likely to bind the membrane-distal dCache_1 domain, while fumarate is more likely to bind the membrane-proximal dCache_1 domain, but binding to the other binding pockets is also possible.

**Figure 2.**
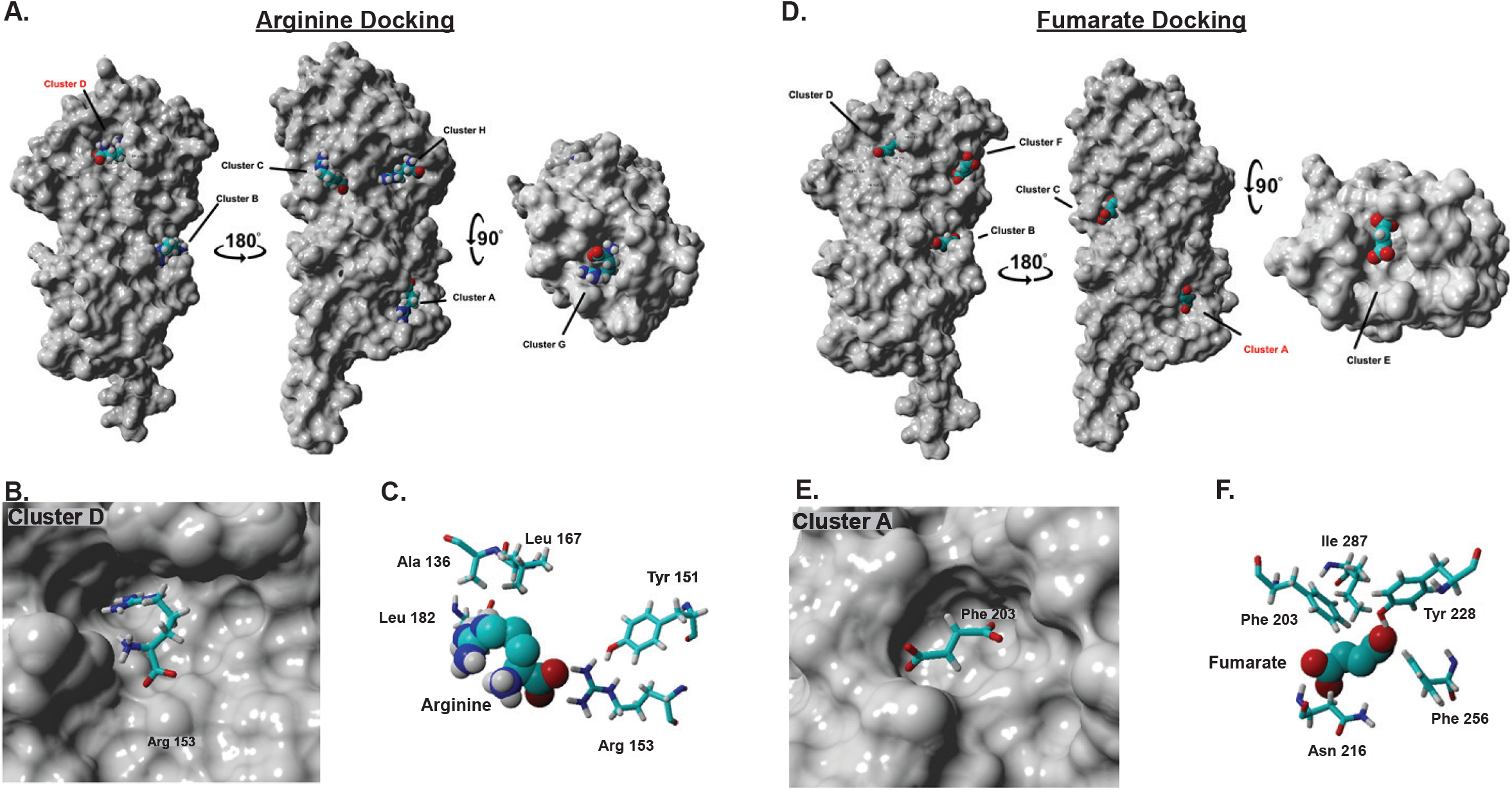
Docking analysis of TlpA and arginine or fumarate identifies several clusters that are occupied by these ligands. Docking analysis shows several clusters occupied by **(A)** arginine and **(D)** fumarate on the surface of a space-filling version of TlpA_LBD_. Cluster C and G are biologically irrelevant due to the homodimer formation of TlpA, and Cluster B and H are poorly populated (Supplemental Table 3). **(B-C)** View of the energetically preferred bound conformation of arginine in Cluster D, the predicted membrane-distal dCache pocket, with key interacting amino acids shown. **(E-F)** View of the energetically preferred bound conformation of fumarate in Cluster A, the membrane-proximal dCache pocket, with key interacting amino acids shown.

To further study TlpA-ligand interactions, we generated TlpA_LBD_ point mutants at residues in the membrane-distal (D165A, M183A) or membrane-proximal (Y228A, Y252A, D254A) binding pockets and determined the binding affinity of the resultant proteins towards all ligands (Table 2 and Fig. 3A). Mutation of either membrane-distal residue resulted in an ∼10-fold decrease in the binding affinity for arginine, cysteine, fumarate, and glucosamine (Fig. 3B and Table 2). Mutation of the membrane-proximal residue TlpA_Y228A_ also lead to a decrease in binding affinity for arginine, cysteine, and fumarate, but it was only 4-fold. The other mutation in the membrane-proximal site (TlpA_D254A_) only affected fumarate binding (Fig. 3B and Table 2). No proximal pocket residues affected glucosamine binding affinity. Membrane-distal and -proximal mutations resulted in a modest ∼3-4-fold increase in binding affinity toward malic acid and thiamine compared to the WT control. Additionally, no appreciable change in binding affinity towards alpha-ketoglutarate was observed for any point mutant. Overall, TlpA binding interactions by the high-affinity ligands arginine, cysteine, fumarate, and glucosamine are most disrupted by mutations in the membrane-distal dCache_1 domain, but mutations in the membrane-proximal dCache_1 also significantly impair fumarate, and to a lesser extent, arginine and cysteine binding, consistent with the predictions from the docking analysis.

**Table 2.**
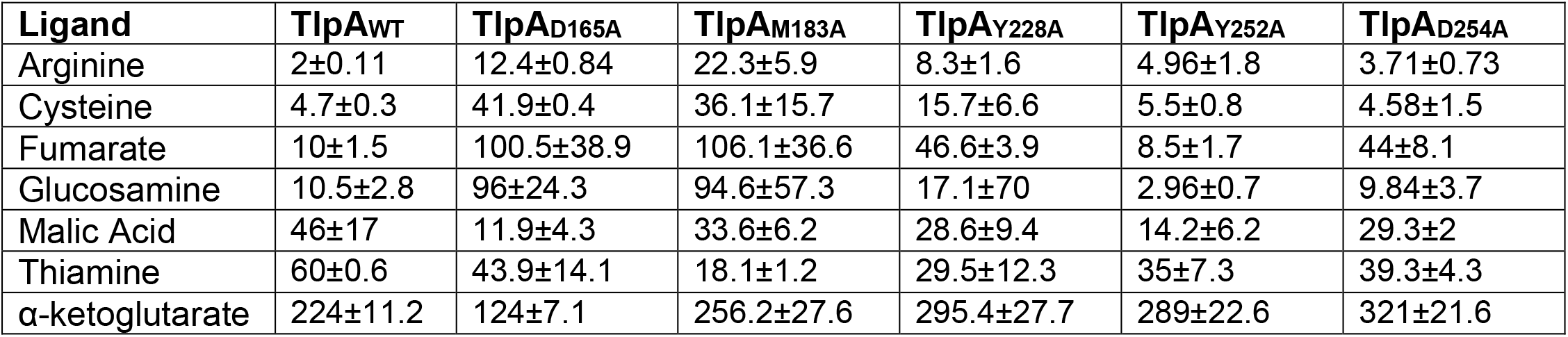
Binding affinity (µM) of TlpA_LBD_ and TlpA_LBD_ membrane-distal and -proximal dCache mutants to TlpA ligands. Data represents the mean values (± SD) of three independent experiments (n= 3).

| Ligand | TlpA<sub>WT</sub> | TlpA<sub>D165A</sub> | TlpA<sub>M183A</sub> | TlpA<sub>Y228A</sub> | TlpA<sub>Y252A</sub> | TlpA<sub>D254A</sub> |
| --- | --- | --- | --- | --- | --- | --- |
| Arginine | 2±0.11 | 12.4±0.84 | 22.3±5.9 | 8.3±1.6 | 4.96±1.8 | 3.71±0.73 |
| Cysteine | 4.7±0.3 | 41.9±0.4 | 36.1±15.7 | 15.7±6.6 | 5.5±0.8 | 4.58±1.5 |
| Fumarate | 10±1.5 | 100.5±38.9 | 106.1±36.6 | 46.6±3.9 | 8.5±1.7 | 44±8.1 |
| Glucosamine | 10.5±2.8 | 96±24.3 | 94.6±57.3 | 17.1±7.0 | 2.96±0.7 | 9.84±3.7 |
| Malic Acid | 46±17 | 11.9±4.3 | 33.6±6.2 | 28.6±9.4 | 14.2±6.2 | 29.3±2 |
| Thiamine | 60±0.6 | 43.9±14.1 | 18.1±1.2 | 29.5±12.3 | 35±7.3 | 39.3±4.3 |
| α-ketoglutarate | 224±11.2 | 124±7.1 | 256.2±27.6 | 295.4±27.7 | 289±22.6 | 321±21.6 |

**Figure 3.**
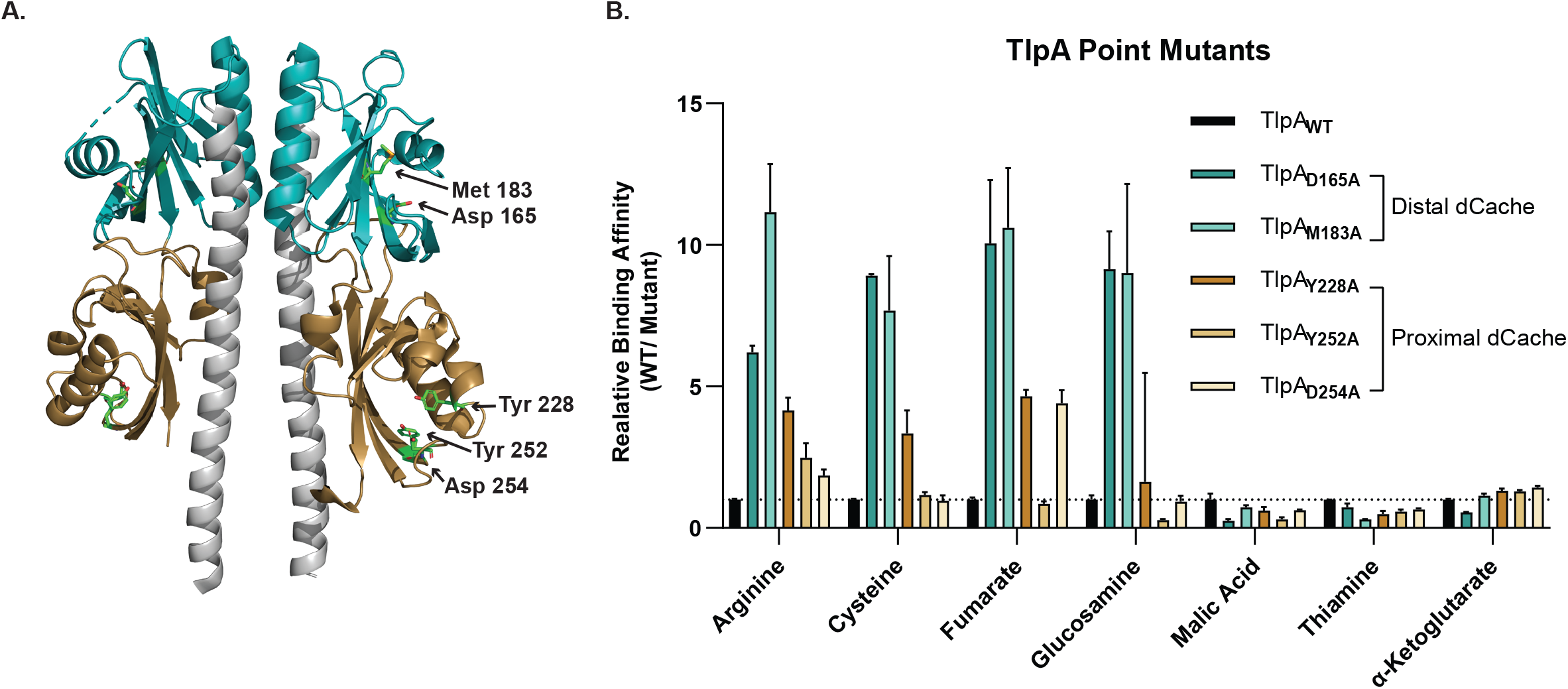
TlpA_LBD_ binds chemotaxis-active ligands through the membrane-distal or -proximal dCache subdomains. **(A)** A ribbon diagram of TlpA_LBD_ as a homodimer is shown. The membrane-distal and membrane-proximal dCache domains are colored teal and gold, respectively. Residues that were mutated to alanine are highlighted in each region to make TlpA_D165A_, TlpA_M183A_, TlpA_Y228A_, TlpA_Y252A_, and TlpA_D254A_. **(B)** The relative binding affinity for each ligand to WT and TlpA_LBD_ membrane-distal and -proximal dCache point mutants. E.g., mutation of D165 to A resulted in a ∼ 6-fold decrease in arginine binding affinity compared to WT binding. Data represented the mean values and standard error of the mean of three independent experiments (n=3).

### TlpA_LBD_ binds arginine and fumarate through distinct binding sites

The above data suggest that TlpA_LBD_ can bind ligands in both dCache_1 subdomain binding pockets. To further analyze the possibility of two distinct binding sites for chemotaxis-active ligands in TlpA_LBD_, we employed a competition SPR (A-B-A) binding assay focused on arginine and fumarate because they were both chemoattractants and predicted to bind different sites preferentially. In this assay, the competition for binding to TlpA_LBD_ between arginine and fumarate is assessed by adding the ligands sequentially and monitoring whether the SPR signal changes upon addition of the second ligand. The two ligands’ binding status can be classified as either independent, shared, or preferential shared sites. For independent sites, ligand A saturates all its binding sites, and ligand B then binds to its independent site; this mode produces additive effects on the SPR signal. Shared sites, in contrast, do not produce additive/cumulative effects, *i*.*e*., ligand A binds its site and then blocks ligand B from the same site. Lastly, it is also possible to have preferential shared sites where ligands share the same binding site, but the protein binds to one ligand preferentially when in equilibrium.

We first saturated TlpA_LBD_ with arginine and then added fumarate. In this case, an increased response (additive effect) was observed, compared to the theoretical value (Fig. 4). This outcome suggests fumarate and arginine bind to independent sites. Conversely, when TlpA_LBD_ was saturated with fumarate, arginine did not produce an additional response, compared to the theoretical value (Fig. 4). This result suggests that fumarate prevented arginine binding, either because arginine competed with fumarate at the same site(s), or that fumarate caused an allosteric effect that prevents arginine binding, a common occurrence in sensory proteins (Biemann and Koshland, 1994). Overall, the docking and competitive SPR data support the hypothesis that there are two binding sites with possible cooperative interactions or overlap between them.

**Figure 4.**
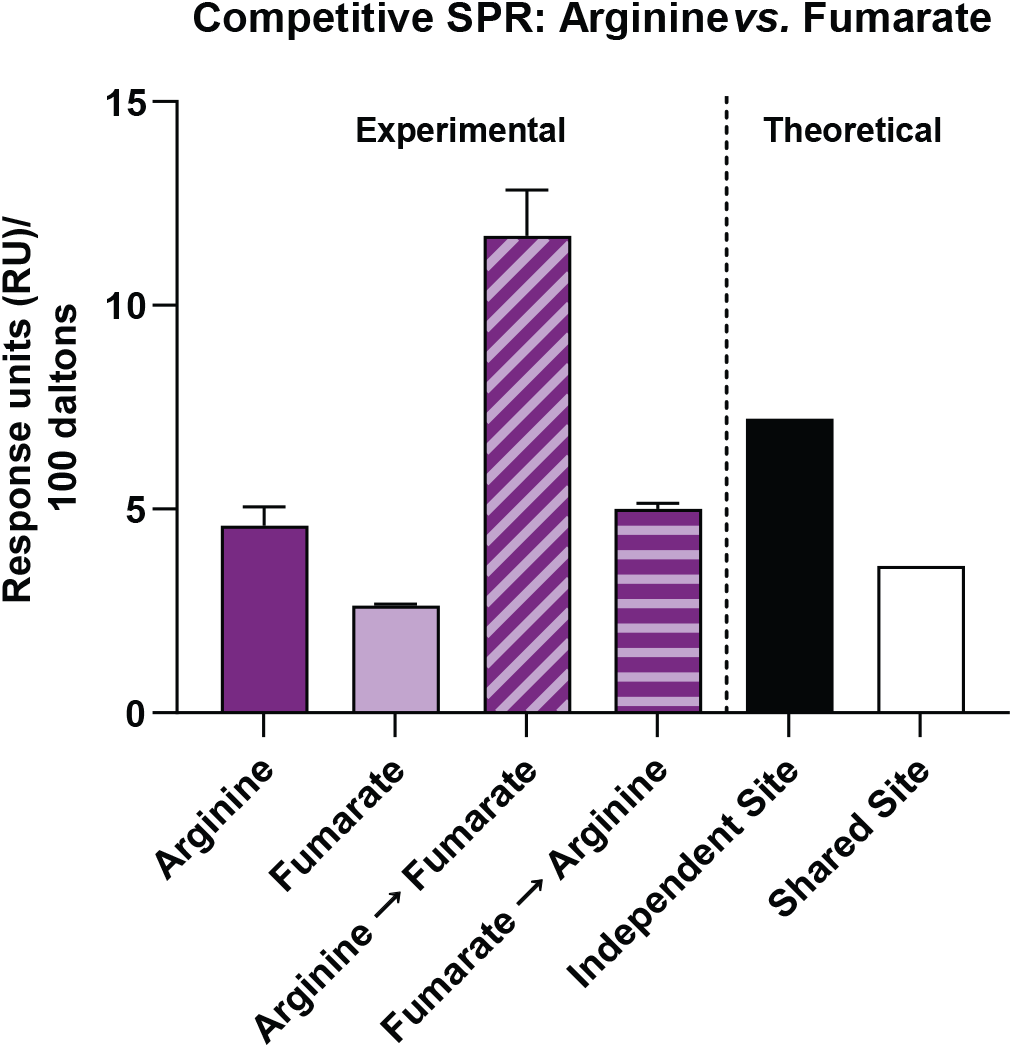
SPR competition analysis indicates the presence of two distinct binding sites for TlpA ligands. SPR competition analysis of binding by arginine and fumarate to WT TlpA_LBD_. Compounds were used at concentrations 10x their respective Kd. Arginine, response to arginine only; Fumarate, response to fumarate only; Arginine→Fumarate, fumarate response after saturation with arginine; and Fumarate→ Arginine, arginine response after saturation with fumarate. The theoretical value is responses units (RU) values based on mathematical theory: Independent Site is the sum of individual responses; Shared Site is the sum of individual responses divided by the number of individual responses. All response data was normalized 100 Da molecular weight for each analyte allowing direct comparison of responses.

The docking analysis and competition SPR assay suggested that arginine and fumarate bind to distinct TlpA_LBD_ sites, therefore we sought to further characterize TlpA-ligand interactions using Saturation Transfer Difference (STD) NMR spectroscopy, which can measure protein-ligand interactions and ascertain which part of a ligand interacts with the receptor protein (Haselhorst et al., 2009; Mayer and Meyer, 1999). When TlpA_LBD_ bound fumarate, a significant STD NMR signal was detected, consistent with the two ethylene protons interacting with the protein (Fig. 5A). Similarly, when TlpA_LBD_ bound arginine, significant STD NMR signals were observed (Fig. 5B). On arginine, the relative STD NMR effects showed the H-3 and H-4 protons on the side chain received the largest saturation transfer from the protein protons, indicating arginine interacts with TlpA_LBD_ around its middle carbon side chain region (Fig. 5B). These results, therefore, provide additional confirmation that TlpA interacts with fumarate and arginine, mostly along the carbon chain backbones in each ligand.

**Figure 5.**
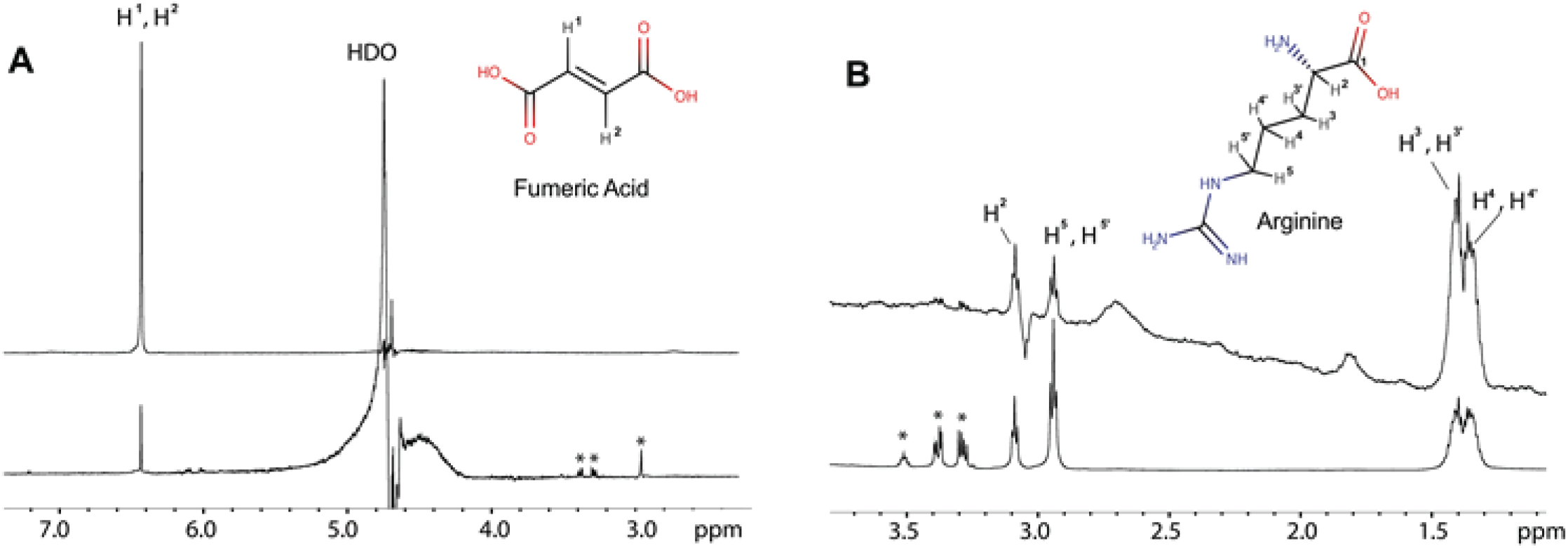
STD NMR analysis supports TlpA_LBD_ binds fumarate and arginine. ^1^H NMR spectra are shown at the bottom for **(A)** fumarate and **(B)** arginine. The STD NMR spectra are shown on top acquired at 600 MHz, 289 K with an on-resonance of -1 ppm and off-resonance of 33 ppm and a total saturation time of 2 sec.

### A non-chemotaxis active TlpA ligand can antagonize chemoattractant responses

It was surprising to find a high affinity direct binding ligand, glucosamine, which bound to the membrane-distal dCache_1 subdomain (Fig. 3) that did not elicit a chemotaxis response (Fig. 1D). Previous reports on ligand interactions with chemoreceptors in *E. coli* and *P. aeruginosa* suggested that some ligands bind chemoreceptors as antagonists, blocking normal chemotactic responses toward chemotaxis-active ligands (Bi et al., 2013; Martín-Mora et al., 2018). Consequently, we tested whether glucosamine could block binding of the chemotaxis-active TlpA ligands arginine and fumarate, using a competitive SPR assay. The results showed that when glucosamine was added following saturation with arginine or fumarate, the response was not additive, suggesting glucosamine competed with both arginine and fumarate (Fig. 6A and B). However, when arginine or fumarate were added to TlpA_LBD_ following initial saturation with glucosamine no additive response was observed (Fig. 6A and B). This result suggests that glucosamine can prevent binding of both ligands to TlpA_LBD_.

**Figure 6.**
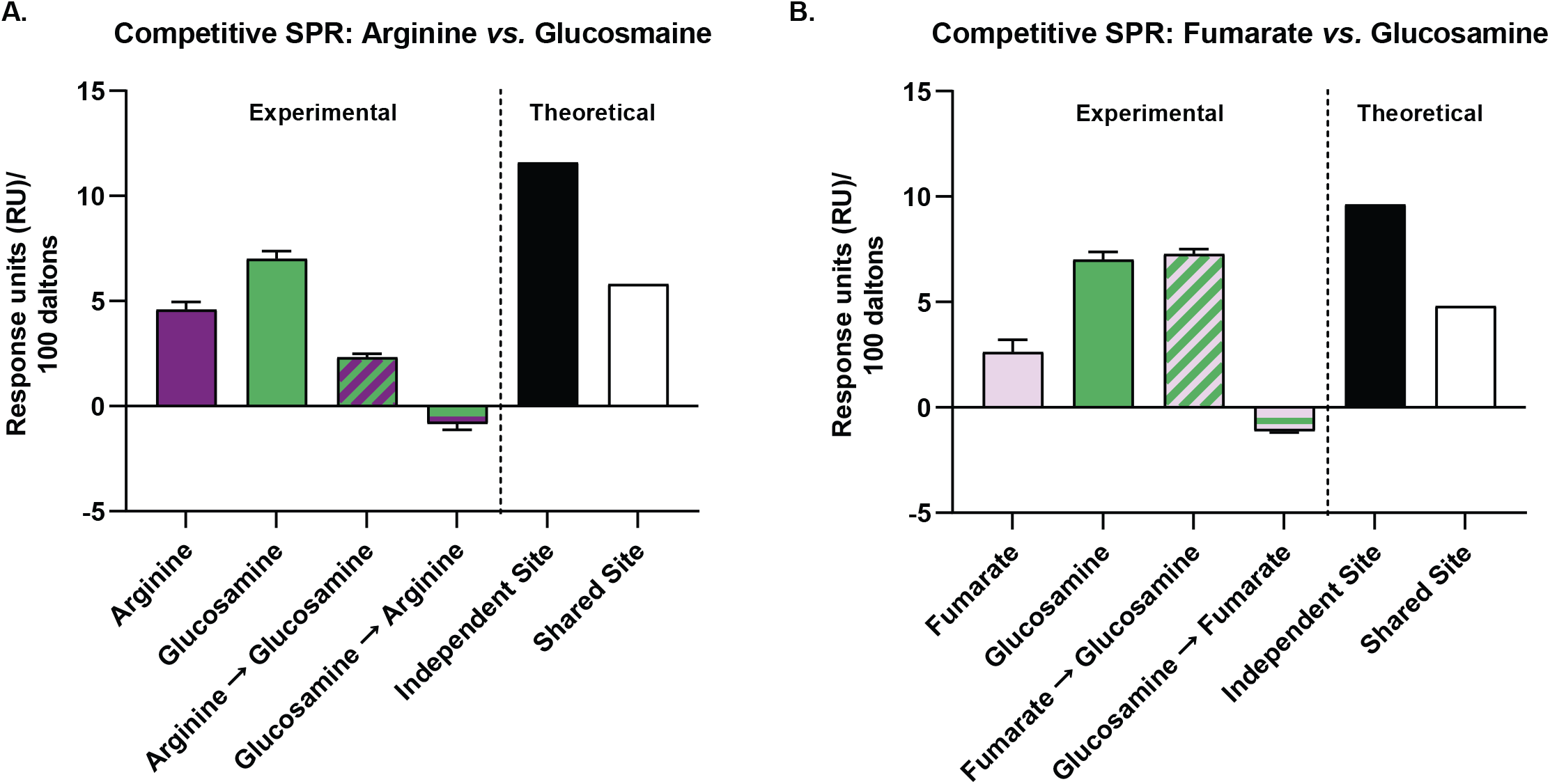
SPR competition analysis demonstrates glucosamine blocks bindings of TlpA chemoattractants. Data from SPR competition analysis of binding of arginine, fumarate, and glucosamine to WT TlpA_LBD_ are shown. Compounds were used at concentration 10x their respective Kd’s. **(A)** Arginine, response to arginine only; Glucosamine, response to glucosamine only; Arginine→Glucosamine, response to glucosamine following saturation with arginine; Glucosamine→Arginine, response to arginine following saturation with glucosamine. **(B)** Fumarate, response to fumarate only; Glucosamine, response to glucosamine only ; Fumarate→Glucosamine, response to glucosamine following saturation with fumarate; Glucosamine→Fumarate, response to fumarate following saturation with glucosamine. The theoretical values are response units (RU) values based on mathematical theory. All response data was normalized 100 Da molecular weight for each analyte allowing direct comparison of responses.

Consequently, we tested whether glucosamine affected *H. pylori* chemotaxis by developing a ligand competition tracking assay between non-chemotaxis active and chemotaxis-active TlpA ligands. This assay is a modified version of our live-cell video microscopy assay where the addition of a chemotaxis-active ligand is followed by the addition of a non-chemotaxis active ligand 10-seconds later and vice versa. Using this approach, we determined that glucosamine addition prevented the chemoattractant response towards arginine, fumarate, and severely blunted the response to cysteine (Fig. 7). This response was decreased regardless of whether glucosamine was added prior to or after the addition of the chemoattractant. In total, these results suggest that glucosamine blocks chemotaxis-active ligand binding and acts as a TlpA chemotaxis antagonist.

**Figure 7.**
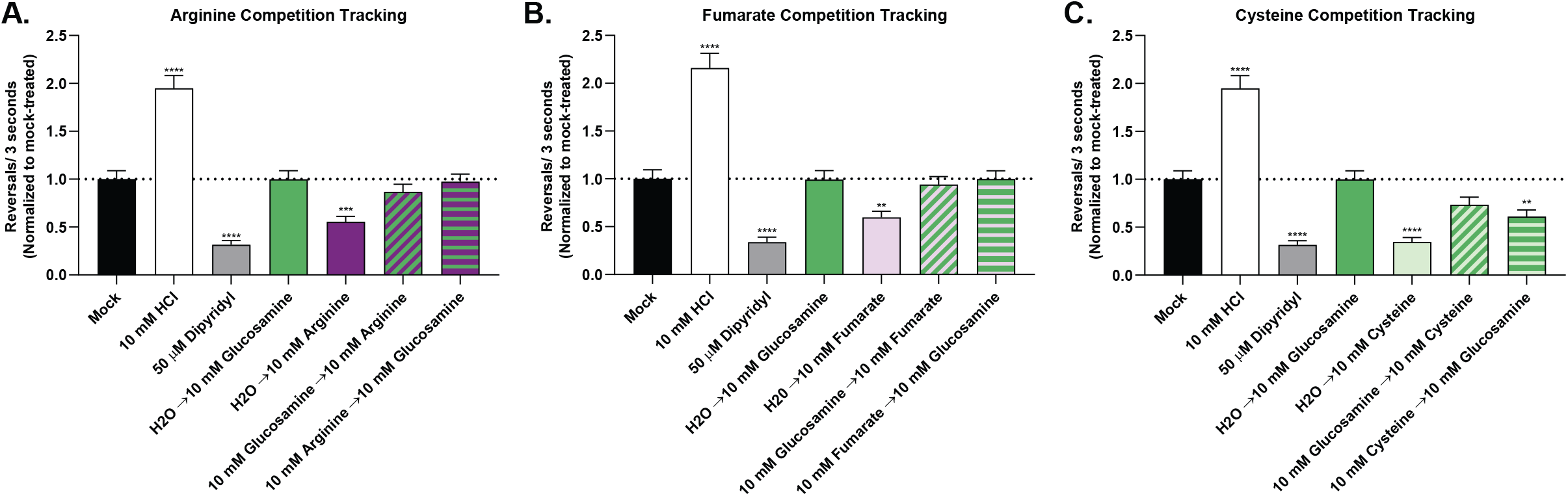
Ligand competition tracking experiment between chemoactive and non-chemoactive TlpA ligands. Cultures of *H. pylori* PMSS1 WT were grown in BB10 overnight and then back-diluted as described in Fig. 1. Cultures were mock-treated or with various concentrations of compounds as indicated. The pH of the cysteine stock was adjusted using NaOH to match the pH of the water used for the untreated control. The cells were immediately filmed, and direction changes were counted over a 3 second swimming period in at least 100 cells per treatment from 3 biological replicates. Repellents increase in direction changes, as exemplified by the control repellent HCl, while attractants decrease direction changes as exemplified by the control attractant dipyridyl. Data is normalized to the untreated control for each strain, as described in the methods. Error bars represent the standard error of the mean. *, P<0.05; **, P<0.01; ***, P<0.001, comparisons to untreated control per strain using two-way ANOVA, Dunnett’s multiple comparison test.

## Discussion

We report the identification of several ligands specific for *H. pylori* dCache_1 chemoreceptor TlpA, including confirmation of previous reports that TlpA interacts with arginine (Cerda et al., 2003, 2011). Arginine, along with fumarate and cysteine, functioned as TlpA-sensed chemotaxis attractants. Furthermore, these chemotaxis-active ligands appeared to interact with the membrane-distal and -proximal dCache_1 domains. Finally, we found that glucosamine acts as a chemotaxis antagonist, blocking TlpA binding and responses to multiple attractants.

### TlpA can bind a broad set of ligands with diverse biological functions

TlpA bound a broad set of molecules ranging from the amino acid arginine with a large, charged side chain, the amino acid cysteine with a smaller polar side chain, organic acids including fumarate, malic acid, and alpha-ketoglutarate, the large vitamin thiamine, and the amino sugar glucosamine. Ultimately, our data suggest that only some of these ligands bind within the canonical dCache_1 binding pockets, including all those that affected chemotaxis. Thus, these results agree with previous reports showing that individual dCache chemoreceptors can sense diverse types of ligands (Rahman et al., 2014; Ud-Din et al., 2020).

The three molecules that elicited a chemotaxis response, arginine, fumarate, and cysteine, have been shown to have important biological roles in *H. pylori* biology. Arginine is an essential amino acid for *H. pylori* under *in vitro* growth conditions (Nedenskov, 1994), and *H. pylori* uses arginine to promote acid tolerance and dampen host immune responses (Alam et al., 2018; Gobert and Wilson, 2016; Mcgee et al., 1999; Valenzuela et al., 2014). Fumarate is predicted to be an alternative terminal electron acceptor for growth under anaerobic respiration (Mendz et al., 1995), and the associated enzyme, fumarate reductase, is essential for *H. pylori* colonization *in vivo* (Ge et al., 2000). Additionally, fumarate is highly depleted when *H. pylori* is co-cultured with gastric organoids, consistent with the prediction that it is preferentially used *in vivo* (Keilberg et al., 2020). Lastly, cysteine is an essential amino acid for some strains of *H. pylori* (Nedenskov, 1994). Each chemotaxis-active TlpA ligand is important for critical cellular functions for *H. pylori*, therefore the ability to sense these ligands a likely survival-linked evolutionary adaptation.

### TlpA appears to use both binding pockets

dCache_1 chemoreceptors contain two potential ligand binding pockets in each Cache subdomain. Previous work showed dCache_1 chemoreceptors sense chemotaxis-active ligands through either subdomain, but as of yet, not both (Machuca et al., 2017; Mckellar et al., 2015). In contrast, our data suggests that TlpA might bind ligands in both Cache subdomains. Ligand binding locations, suggested by docking prediction analysis, placed fumarate within the membrane-proximal subdomain and arginine within the membrane-distal subdomain. These location assignments were further supported by TlpA_LBD_ point mutants of residues within the predicted dCache_1 membrane-distal (D165A and M183A) and membrane-proximal (Y228A, Y252A, D254A) subdomains. All membrane-distal subdomain mutations led to a decreased binding affinity for the chemotaxis-active ligands, arginine, fumarate, cysteine, as well as the antagonist ligand glucosamine. These findings suggest that the membrane-distal site is important for chemotaxis signaling, as seen in other dCache_1 receptors (Mckellar et al., 2015; Upadhyay et al., 2016). It has been shown that individual dCache_1 receptors can bind aliphatic, small polar, and large, positively charged amino acids through a single subdomain due to the malleable nature of dCache_1 receptors that can accommodate ligands of different sizes and charge (Ud-Din et al., 2020). Thus, it is plausible the membrane-distal subdomain could be able to accommodate these diverse ligands.

Our data also suggest that the membrane-proximal subdomain plays a role in the TlpA ligand binding. Site-directed mutagenesis of the membrane-proximal subdomain, as well as the membrane-distal one, decreased fumarate binding. This outcome suggests that the point mutations either directly disrupted ligand binding or altered long-range interactions in the protein that influence ligand binding affinities. Additionally, our data showed that fumarate blocked arginine binding, despite having similar binding affinities. Several models could account for these findings. One is that fumarate binds at the membrane-proximal site and creates an allosteric change that prevents ligand binding at the membrane-distal site. Alternatively, the amino acid changes in the proximal site could affect ligand affinities at the distal site. Finally, a third possibility is that arginine binds at both sites, and proximal site binding is affected by the proximal mutations. It will be interesting to dissect whether there is cooperative interactions, distinct site binding, or simultaneous site binding. While cooperativity between subdomains in dCache_1 chemoreceptors has not yet been observed, it is well documented that four-helix bundle types of chemoreceptors have negative cooperativity between their two binding sites (Biemann and Koshland, 1994). It furthermore is not yet known which site is required for chemotaxis. Overall, our data suggests that TlpA may use both dCache_1 subdomains to bind ligands.

### TlpA chemotaxis responses can be antagonized

We were somewhat surprised to find a high-affinity binding TlpA ligand, glucosamine, that did not elicit a chemotaxis response yet appeared to bind the membrane-distal dCache_1 subdomain. Indeed, we found that glucosamine occluded chemotaxis-active TlpA ligands from binding and inhibited the normal chemoattractant responses towards arginine, fumarate, and cysteine. This response was observed whether glucosamine was added before or after the addition of the TlpA chemoattractants, suggesting glucosamine may have a very high on-rate for binding TlpA compared to arginine, fumarate, or cysteine. Two studies have reported high-affinity chemoreceptor ligands that acted as antagonists by blocking chemotaxis-active ligand binding (Bi et al., 2013; Martín-Mora et al., 2018), and Cache receptor antagonists have been reported for a histidine kinase (Busch et al., 2007). Martín-Mora and colleagues showed that binding of the attractant malic acid to the *P. aeruginosa* sCache chemoreceptor PA2652 was inhibited by either citraconic acid or D,L-methylsuccinic acid, and subsequently chemoattractant responses towards malic acid were decreased (Martín-Mora et al., 2018). Another of these studies described finding diverse ligands for the *E. coli* four-helix bundle chemoreceptor Tar, and reported a high-affinity ligand, *cis*-1,2-cyclohexane-dicarboxylic acid also acted as an antagonist for aspartate chemotaxis. *cis*-1,2-cyclohexane-dicarboxylic acid competed for aspartate binding and blocked intracellular kinase activity (Bi et al., 2013). We expect there will be more discoveries of these types of chemo-modulatory antagonists, or maybe even chemotaxis enhancing ligands, because ligand discovery methods have changed. Specifically, previous efforts relied on chemotaxis assays, and so only chemotaxis-active ligands could be identified. In contrast, recent approaches look for direct ligand-receptor interactions at the molecular level, thus expanding our ability to identify the interacting partners (Bi et al., 2013; Ehrhardt et al., 2018; Hartley-Tassell et al., 2010; Krell, 2015; Mckellar et al., 2015).

The function of chemoreceptor antagonists is not yet known in any system (Bi et al., 2013; Martín-Mora et al., 2018). In the case of TlpA, it is possible to speculate that when confronted with abundant glucosamine, *H. pylori* benefits by not responding to arginine, fumarate, or cysteine. However, the role of glucosamine in *H. pylori* infection is unknown. Chemotaxis antagonists may be useful tools to modulate chemotaxis and affect bacterial pathogenesis. In the case of TlpA, blocking its function early in infection would decrease colonization; however, a later attenuation of chemotactic responses might be predicted to enhance inflammation(Andermann et al., 2002; Rolig et al., 2012; Williams et al., 2007).

Expanding the knowledge of TlpA ligands, and how TlpA interacts with these ligands, is essential for better understanding why TlpA enhances the *in vivo* fitness of *H. pylori* and alters inflammatory phenotypes driven by *H. pylori*. Future experiments manipulating the ability of *H. pylori* to sense specific TlpA ligands will be useful to understand whether all or a subset of TlpA ligands play a role in driving these *in vivo* phenotypes (Andermann et al., 2002; Rolig et al., 2012; Williams et al., 2007). Furthermore, this work provides another example (Bi et al., 2013; Martín-Mora et al., 2018) of a chemotaxis system having antagonistic ligands, operating through a distinct type of chemoreceptor ligand-binding domain. These results suggest an interesting possible mechanism for regulating responses to multiple chemotactic ligands in a nutrient-rich environment using agonist and antagonist ligands.

## Methods

### TlpA construct design and protein purification

The periplasmic portion of TlpA (TlpA_LBD_), amino acids 28-299, from *Helicobacter pylori* SS1 was cloned into a pBH4 expression vector (pBH4_TlpA_LBD_) to generate an N-terminal 6x-His-tagged construct, with a TEV protease site, under control of IPTG (Sweeney et al., 2018). Alanine point mutants at Asp 165, Met 183, Tyr 228, Tyr 252, and Tyr 254 in TlpA_LBD_ were generated via site-directed mutagenesis of pBH4_TlpA_LBD_ with primers listed in Supplemental Table 4 (QuikChange, Stratagene). TlpA_LBD_ and all point mutants were purified as described by Sweeney *et al*. 2018(Sweeney et al., 2018). CD spectroscopy was used to confirm the correct folding of all proteins (Supplemental Fig. 3) (Elgamoudi et al., 2021).

### Ligand binding array

Small-molecule arrays were prepared and performed as previously described (Day and Korolik, 2018). Briefly, 1 µg of purified TlpA_LBD_ in PBS, pH 7.2, was incubated with an equal molar concentration of anti-His antibody (Cell Signaling), followed by incubation with a secondary antibody (Thermo Scientific) for signal amplification. The protein-antibody mix was added to an Arrayit Superepoxy III glass substrate array blocked with PBS, pH 7.2, with 1% bovine serum albumin. The glass substrate array was printed with quadruplicate spots of amino acids, salts of organic acids, and other small molecules (Supplementary Table 2). Unbound protein was washed away with PBS with 0.05% Tween. The arrays were scanned by a ProScan Array scanner at 488/520 nm and the results analyzed by ScanArray Express software program. Binding was defined as a value greater than 1-fold increase above mean background relative fluorescence units (RFU). The mean background was calculated from the average background of empty spots on the array plus three standard deviations. Statistical analysis of the data was performed by a Student’s t-test with a confidence level of 99.99% (p ≤ 0.0001). All arrays were performed in triplicate with a total of 12 data points for each small molecule tested.

### Surface Plasmon Resonance Measurements

Purified TlpA_LBD_ was immobilized on a CM5 series S sensor chip and binding affinities tested using a BIAcoreTM S200 instrument (GE Healthcare) as described (Day et al., 2016; Day and Korolik, 2018; Rahman et al., 2014). Briefly, proteins were captured using an amine coupling kit (GE Healthcare), in which the carboxylmethyl dextran matrix of the sensor chip was activated by injection of a mixture of 0.2M 1-ethyl-3-[(3-dimethylamino) propyl]-carbodiimide (EDC) and 0.05 M N-hydroxysuccinimide (NHS), followed by neutralization of the remaining unreacted NHS ester groups by an injection of 1 M ethanolamine-HCl (pH 8.0). Purified TlpA_LBD_ was diluted in 10 mM sodium acetate buffer pH 4.5 at a concentration of 100 μg/mL for immobilization to the chip. 8400 response units (RU) of TlpA_LBD_ were captured on Flow Cell 2. As a negative control, Flow Cell 1 was a blank control undergoing the same treatment as the other flow paths, without the protein injection. This set enabled double reference subtraction of the responses (2-1, 3-1, 4-1). The tested compounds were prepared as a stock concentration of 100-200mM in PBS. The compounds were then diluted between 1 nM-1mM in a series of 1:10 dilutions in PBS and run over the flow cells at a flow rate of 30 μL/min. Between each sample testing, a series of buffer-only injections were run to enable double blank subtraction for the sensorgram assessment. After the initial run, based on the results, the dilution series ranged from 0.195 µM to 1 mM in 1:4 dilutions in PBS. Then the samples were run using single-cycle kinetic/affinity methods in triplicate for those compounds that showed sub-millimolar affinity after the initial binding screen. The datasets were analyzed using the BIAcore s200 evaluation software 2.0.2; sensorgrams were double reference subtracted.

### Bacterial strains and growth conditions

For all chemotaxis assays, *H. pylori* strain PMSS1 was used(Arnold et al., 2011). Bacteria were grown in Brucella broth (BD BBL/Fisher) with 10% heat-inactivated fetal bovine serum (FBS) (Life Technologies) (BB10), with shaking, at 37°C, under microaerobic conditions of 5% O_2_, 10% CO_2_, balance N_2_. The PMSS1 *ΔtlpA* mutant was created by natural transformation of wild-type PMSS1 with 5 μg of *ΔtlpA::cat* SS1 genomic DNA(Andermann et al., 2002). Chloramphenicol-resistant mutants were selected using 10 μg/ml chloramphenicol on Columbia Horse Blood Agar as previously described(Andermann et al., 2002). Mutation of *tlpA* was confirmed by PCR amplification of genomic DNA from WT PMSS1, *ΔtlpA::cat* PMSS1, and *ΔtlpA::cat* SS1 using primers TlpA_SS1_5’ (TTGTCTAAAGGTTTGAGTATC) and TlpA_SS1_3’(TTAAAACTGCTTTTTATTCAC) (This study) (Supplemental Figure 4).

### Chemotaxis assays

Swimming behavior assays were done with *H. pylori* PMSS1 strains and grown in BB10 as described above. Overnight cultures were diluted to an OD_600_ of 0.1 in fresh BB10 and then incubated with shaking as above until an OD_600_ of 0.12– 0.15 was reached. Motility of these cultures was confirmed, and then they were used for chemotaxis assays by treating with L-arginine monohydrochloride (J.T. Baker, B577-05), sodium fumarate (Acros Organics, 215531000), L-cysteine hydrochloride monohydrate (RPI, C81020), D(+)-glucosamine hydrochloride (Chem-Impex International Inc., 01450), thiamine hydrochloride (Fisher BioReagents, BP892), α-ketoglutaric acid (Santa Cruz Biotechnology, SC-208504), or L-malic acid (MP Biomedicals, 102237) at a final concentration of 0.1 mM, 1 mM, 10 mM or an equal volume of H_2_O as a mock-treated control (4 μl H_2_O or 4 μl of ligand stock in H_2_O into 96 μl culture). The number of direction changes in a bacterial swimming trajectory were enumerated over a three-second interval to determine whether each putative ligand is sensed as an attractant, repellent, or elicits no response (Collins et al., 2016; Goers Sweeney et al., 2012; Lertsethtakarn et al., 2015; Machuca et al., 2017; Rader et al., 2011; Schweinitzer et al., 2008; Terry et al., 2006). The results were compared to both a repellent control, 10 mM HCl (Fisher Chemical, A144S), which results in increased direction changes (Croxen et al., 2006; Goers Sweeney et al., 2012) and an attractant control, 50 μM 2,2’-dipyridyl (Arcos Organics, 117500250), which results in fewer direction changes (Collins et al., 2016). Each control is sensed by chemoreceptors other than TlpA (Collins et al., 2016; Croxen et al., 2006; Goers Sweeney et al., 2012; Huang et al., 2017). The pH of BB10 upon treatment was independently assessed using a Denver Instruments pH meter. For tracking with ligands that acidified the pH of BB10 upon treatment, the ligand pH was adjusted to the pH of water using NaOH. Cultures were filmed immediately after ligand addition at 400x magnification using a Hamamatsu Digital Camera C4742-95 with the μManager software (Version 1.4.22), mounted on a Nikon Eclipse E600 phase-contrast microscope. For the competition chemotaxis assay, cultures of *H. pylori* and ligands were prepared as above. However, 1 minute after the addition of a non-chemoactive ligand at a final concentration of 10 mM, a chemoactive ligand was added at a final concentration of 10 mM, and then cultures were filmed as described above. Videos were relabeled to blind the observer to the strain identity. For each sample, >100 3-s-long bacterial tracks from three independent cultures were analyzed manually to identify stops followed by direction changes. Data for all biological replicates for each condition was combined and the average number of directions changes in 3 seconds and standard error of the mean were calculated. For each strain, data was normalized to the untreated control for each experimental condition. Statistical analysis of the data for treated versus untreated samples was performed using a two-way ANOVA, Dunnett’s multiple comparison test.

### SPR TlpA competition assays

SPR competition assays were performed by using a BIAcore S200 instrument and the A-B-A inject function(Elgamoudi et al., 2021). Competition A-B-A analyses were used to interrogate the specificity of the potential ligand binding site preferences of TlpA_LBD_ and to unravel the nature of the ligand-sensor interactions. This assay was designed to show if a cumulative response is observed when a second analyte (B) is flown across the bound protein saturated with the first analyte (A) (Fig 4). As the assay is designed to provide saturation of all analytes tested, this assay does not provide 1:1 competition to indicate which is the preferred analyte for a binding site. The proteins, wild type TlpA_LBD_ was immobilized as above. The A-B-A was used with combinations of each of the compounds (at concentration 10 × K_D_) and PBS control, with 60-second injections of analyte A to ensure saturation or near-saturation was reached prior to competition with analyte B. The results were analyzed using BIAcore S200 evaluation software using the sensorgram mode, and data was zeroed to baseline before the initial A injection. All response data was normalized 100 Da molecular weight for each analyte, allowing direct comparison of responses. The theoretical values in responses units (RU) values based on mathematical theory calculated by the sum of individual responses (Independent Sites) or the sum of the individual responses divided by the number of individual responses (Shared Sites).

### Saturation Transfer Difference (STD) NMR

In this experiment, the entire TlpA_LBD_ protein was first saturated at the protein resonances, and then excess ligand was added. As the ligand binds and releases from the receptor, saturation transfers from the protein to bound ligand. This transfer appeared as an increase in ligand intensity on epitopes that interacted with the TlpA_LBD_ protein. For STD NMR experiments, samples of 25 µM TlpA_LBD_ in complex with either 2.5 mM Arginine (Arg) or Fumaric Acid (Fum) in 99% D_2_O were prepared. All STD NMR spectra were acquired in Shigemi tubes (Shigemi, USA) with a Bruker 600 MHz Advance spectrometer at 283 K using 1^H^/13^C^/15^N^ gradient cryoprobe equipped with z-gradients. Protein resonances were saturated at -1.0 ppm (on-resonance) and 33 ppm (off-resonance) and a total saturation time of 2 sec. A total of 512 scans per STD NMR experiment were acquired and a WATERGATE sequence was used to suppress the residual HDO signal. A spin-lock filter with 5 kHz strength and a duration of 10 ms was applied to suppress protein background. On- and off-resonance spectra were stored and processed separately, and the final STD NMR spectra were obtained by subtracting the on- and off-resonance spectra. Control STD NMR experiments were performed identically in the absence of protein.

### Docking analysis

To evaluate a potential binding site for Arginine (Arg) and fumeric acid (Fum) to TlpA_LBD_ (PDB code: 6E09). A blind docking experiment was performed using the AutoDock Vina protocol(Trott and Olson, 2009), a high-scoring molecular docking program(Wang et al., 2016), implemented in the YASARA Structure molecular modeling package (Ver. 16.46)(Krieger et al., 2002). The blind docking experiment was set up by using the entire TlpA as-potential binding site (grid size 92.99 Å x 75.73 Å x 62.13 Å). A total number of 999 Vina docking runs were performed.

## Acknowledgments

We like to thank Shaui Hu and Xiaolin Liu for their thoughtful comments on this article. We thank Emily Sweeney and Karen Guillemin for sharing the structure of TlpA (PBD: 6E09) before its publication and for their thoughtful comments on this article. The described project was supported by the National Institute of Allergy and Infectious Diseases (NIAID) grant to KMO. The funders had no role in study design, data collection, and interpretation, or decision to submit the work for publication.

## Notes

### Competing Interest Statement

The authors have declared no competing interest.

