## Supplemental Material for "The immunomodulatory dCache chemoreceptor TlpA of *Helicobacter pylori* binds multiple attractant and antagonistic ligands via distinct sites"

### Supplemental Figure 1

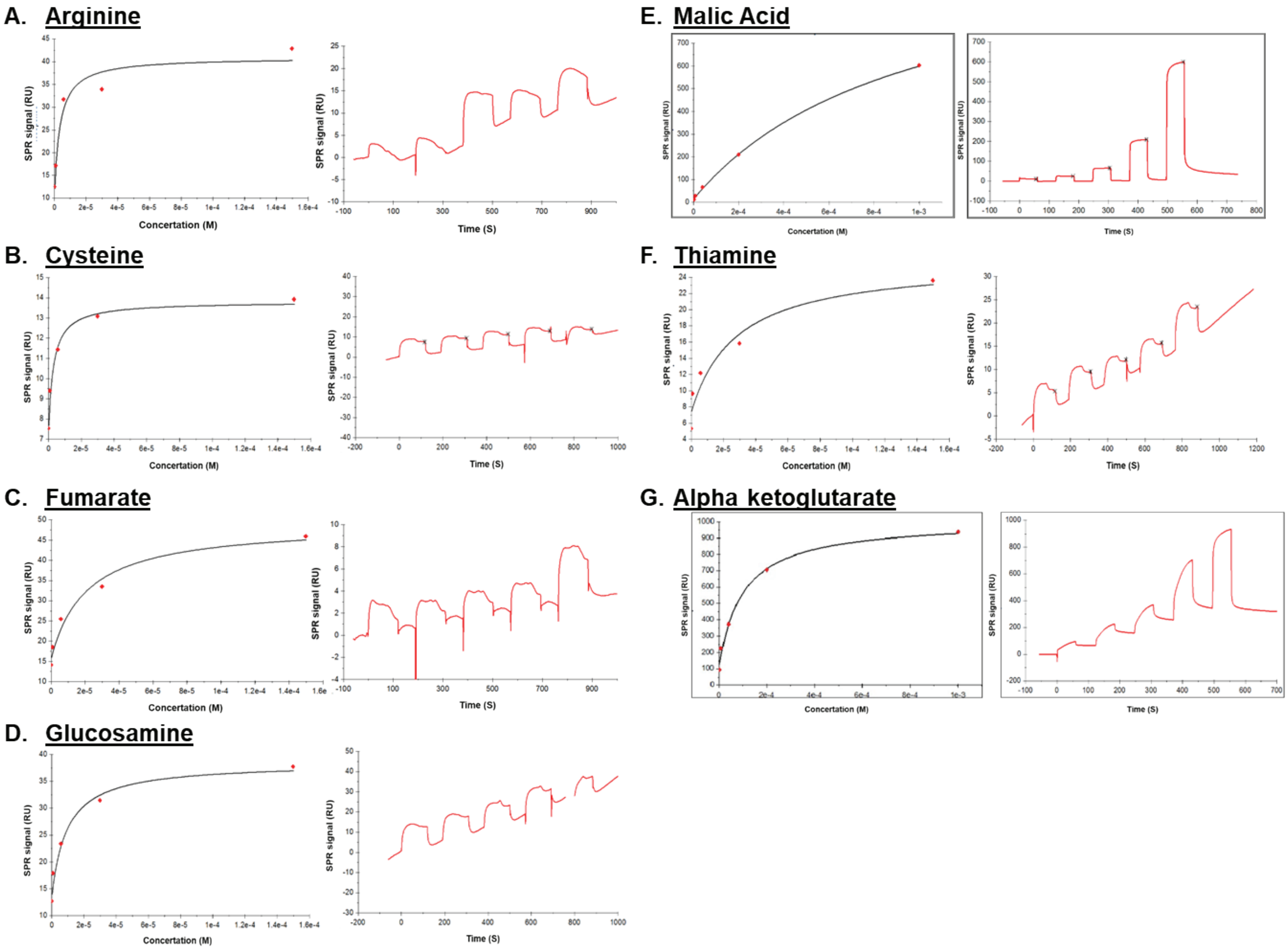


**Supplemental Figure 1- Representative sensorgrams from surface plasmon resonance (SPR) analysis of TlpA$_{LBD}$ with A) arginine, B) Cysteine, C) Fumarate, D) Glucosamine, E) Malic Acid, F) Thiamine, G) Alpha ketoglutarate.**

### Supplemental Figure 2

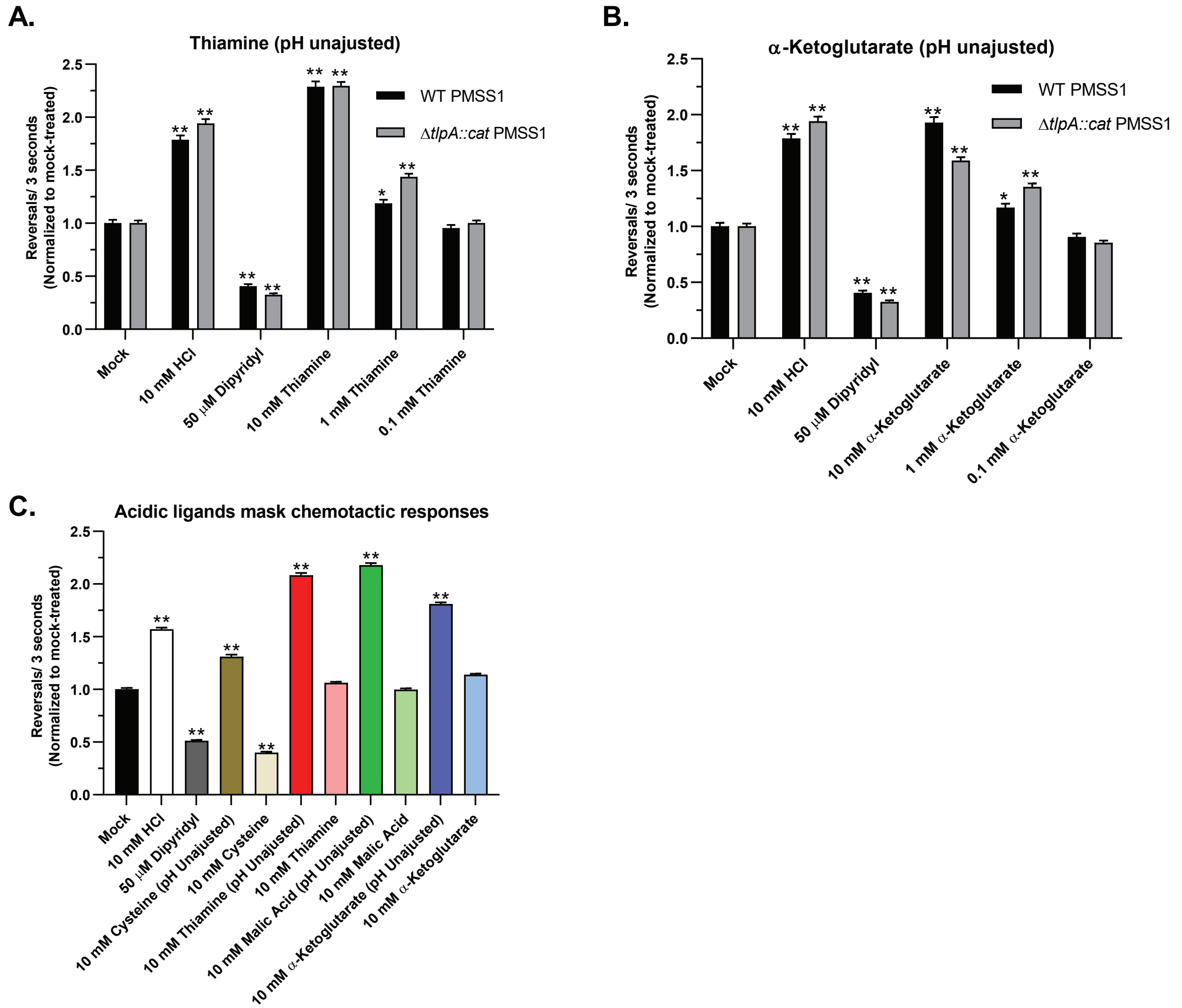


**Supplemental Figure 2- Thiamine, alpha-ketoglutarate, cysteine, and malic acid trigger TlpA-independent repellent responses. (A-B)** Cultures of WT and *ΔtlpA::cat* PMSS1 were grown in BB10 overnight and then treated with various concentrations of thiamine or alpha-ketoglutarate. **(C)** Cultures of WT PMSS1 were grown as above and treated with 10mM pH neutralized or unadjusted thiamine, alpha-ketoglutarate, cysteine, and malic acid. For all experiments, cells were immediately filmed after ligand addition, and direction changes were counted over a 3 second swimming period in at least 100 cells per treatment. Repellents cause increases in direction changes, as exemplified by the control repellent HCl, while attractants cause fewer direction changes as exemplified by the control attractant dipyridyl. Error bars represent the standard error of the mean. *, $P<0.05$; **, $P<0.01$, comparisons to untreated control per strain using two-way ANOVA, Dunnett's multiple comparison test.

**Supplemental Figure 3**

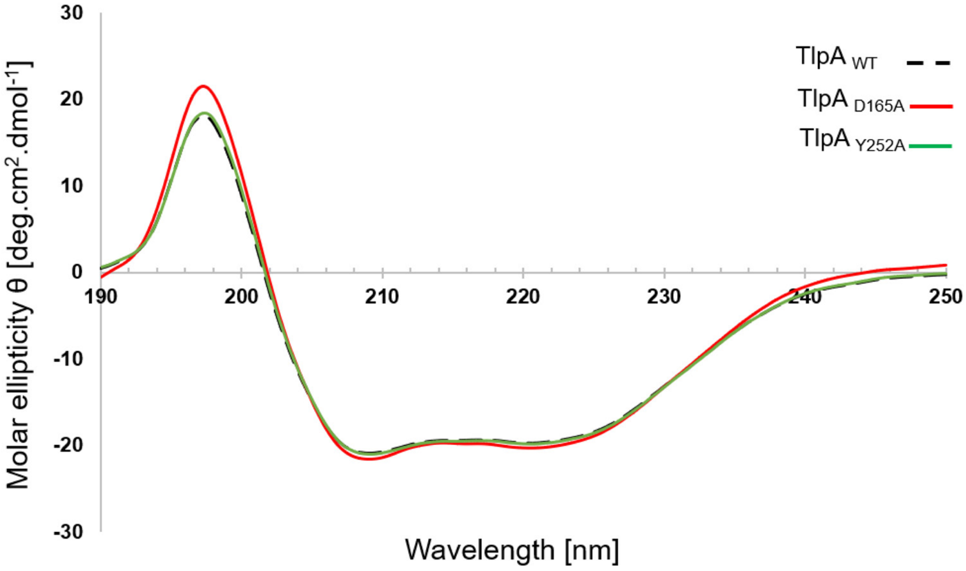


**Supplemental Figure 3- CD spectrometry of WT TlpA$_{LBD}$ and point mutants TlpA$_{D165A}$ and TlpA$_{Y252A}$.** The far-UV CD spectra of TlpA$_{D165A}$ and TlpA$_{Y252A}$ are similar to WT TlpA$_{LBD}$, including all other TlpA$_{LBD}$ point mutants used in this study (Data not shown).

**Supplemental Figure 4**

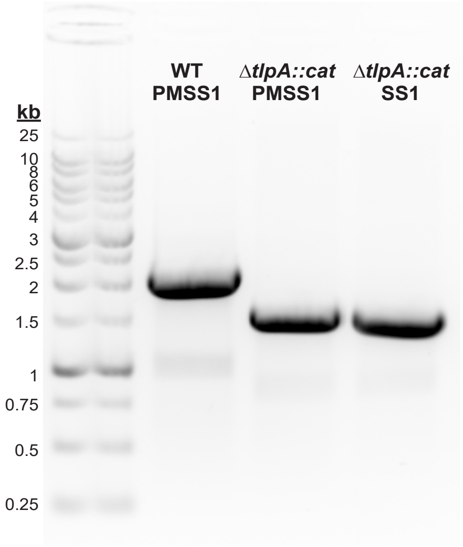


**Supplemental Figure 4- Verification of *ΔtlpA::cat* PMSS1.** Mutation of *tlpA* was confirmed by PCR amplification of gDNA from WT PMSS1, Δ*tlpA::cat* PMSS1, and Δ*tlpA::cat* SS1 (Andermann et al., 2002) using primers TlpA_SS1_5’ (TTGTCTAAAGGTTTGAGTATC) and TlpA_SS1_3’(TTAAAACT-GCTTTTTATTCAC).

**Supplemental Table 1- Amino acids, salts of organic acids, other chemotactic small molecules and glycans printed on the small molecule chemotaxis arrays.**

| Name | Structure/ Molecular Formula |
|---|---|
| **Amino acids, salts of organic acids, + other small molecules** | |
| Alanine | $C_3H_7NO_2$ |
| Arginine | $C_6H_{14}N_4O_2$ |
| Asparagine | $C_4H_8N_2O_3$ |
| Aspartate | $C_4H_7NO_4$ |
| Cysteine | $C_3H_7NO_2S$ |
| Fumaric Acid | $C_4H_4O_4$ |
| Glucosamine | $C_6H_{13}NO_5$ |
| Glutamic acid | $C_5H_9NO_4$ |
| Glutamine | $C_5H_{10}N_2O_3$ |
| Histidine | $C_6H_9N_3O_2$ |
| Isoleucine | $C_6H_{13}NO_2$ |
| leucine | $C_6H_{13}NO_2$ |
| Lysine | $C_6H_{14}N_2O_2$ |
| Malic Acid | $C_4H_6O_5$ |
| Methionine | $C_5H_{11}NO_2S$ |
| Phenylalanine | $C_9H_{11}NO_2$ |
| Proline | $C_5H_9NO_2$ |
| Purine | $C_5H_4N_4$ |
| Serine | $C_3H_7NO_3$ |
| Succinic Acid | $C_4H_6O_4$ |
| Thiamine | $C_{12}H_{17}N_4OS$ |
| Threonine | $C_4H_9NO_3$ |
| Tryptophan | $C_{11}H_{12}N_2O_2$ |
| Tyrosine | $C_9H_{11}NO_3$ |
| Valine | $C_5H_{11}NO_2$ |
| α-ketoglutarate | $C_5H_6O_5$ |
| **Terminal Galactose** | |
| Lacto-*N*-Biose I | Galβ1-3Glc*N*Ac |
| N-Acetyllactosamine | Galβ1-4Glc*N*Ac |
| β1-4galactosyl-galactose | Galβ1-4Gal |
| β1-6galactosyl-*N*-acetylglucosamine | Galβ1-6Glc*N*Ac |
| β1-3galactosyl-*N*-acetylgalactosamine | Galβ1-3Gal*N*Ac |
| Asialo$G_{M1}$ | Galβ1-3Gal*N*Acβ1-4Galβ1-4Glc |
| Lacto-*N*-tetrose | Galβ1-3Glc*N*Acβ1-3Galβ1-4Glc |
| Lacto-*N*-neotetrose | Galβ1-4Glc*N*Acβ1-3Galβ1-4Glc |
| Lacto-*N*-neohexose | Galβ1-4Glc*N*Acβ1-6(Galβ1-4Glc*N*Acβ1-3)Galβ1-4Glc |
| Lacto-*N*-hexose | Galβ1-4Glc*N*Acβ1-6(Galβ1-3Glc*N*Acβ1-3)Galβ1-4Glc |
| Globotriose | Gala 1-4Galβ1-4Glc |

**Cont. Supplemental Table 1**

| | |
|---|---|
| Tn Antigen | Gal*N*Aca1-*O*-Ser |
| Galactosyl-Tn Antigen | Galb1-3Gal*N*Acα1-*O*-Ser |
| α1-3 Galactobiose | Gala1-3Gal |
| Linear B-2 Trisaccharide | Gala1-3Galβ1-4Glc*N*Ac |
| Linear B-6 Trisaccharide | Gala1-3Galβ1-4Glc |
| α1-3, β1-4, α1-3 Galactotetrose | Gala1-3Galβ1-4Galα1-3Gal |
| β1-6Galactobiose | Galβ1-6Gal |
| Terminal disaccharide of globotriose | Gal*N*Acβ1-3Gal |
| Receptor for pili of *P. aeruginosa* | Gal*N*Acβ1-4Gal |
| P1 Antigen | Galα1-4Galβ1-4Glc*N*Ac |
| α-D-*N*-acetylgalactosaminyl-1-3Gal-β1-4Glc | Gal*N*Acα1-3Galβ1-4Glc |
| iso-Lacto-*N*-octose | Galβ1-3Glc*N*Acβ1-3Galβ1-4Glc*N*Acβ1-6(Galβ1-3Glc*N*Acβ1-3)Galβ1-4Glc |
| *para*-Lacto-*N*-hexose | Galβ1-3Glc*N*Acβ1-3Galβ1-4Glc*N*Acβ1-3Galβ1-4Glc |
| **Terminal *N*-Acetyl glucosamine** | |
| *N,N′*-Diacetyl chitobiose | Glc*N*Acβ1-4Glc*N*Ac |
| *N,N′,N″*-Triacetyl chitotriose | Glc*N*Acβ1-4Glc*N*Acβ1-4Glc*N*Ac |
| *N,N′,N″,N‴*-Tetraacetyl chitotetrose | Glc*N*Acβ1-4Glc*N*Acβ1-4Glc*N*Acβ1-4Glc*N*Ac |
| *N,N′,N″,N‴,N″″,N″‴*-Hexaacetyl chitohexose | Glc*N*Acβ1-4Glc*N*Acβ1-4Glc*N*Acβ1-4Glc*N*Acβ1-4Glc*N*Acβ1-4Glc*N*Ac |
| Bacterial cell wall muramyl discaccharide | Glc*N*Acβ1-4Mur*N*Ac |
| **Mannose containing structures** | |
| β1-2-*N*-Acetylglucosamine-mannose | Glc*N*Acβ1-2Man |
| Bianntennary *N*-linked core pentasaccharide | Glc*N*Acβ1-2Manα1-6(Glc*N*Acβ1-2Manα1-3)Man |
| α1-2-Mannobiose | Manα1-2Man |
| α1-3-Mannobiose | Manα1-3Man |
| α1-4-Mannobiose | Manα1-4Man |
| α1-6-Mannobiose | Manα1-6Man |
| α1-3, a1-6-Mannobiose | Manα1-6(Manα1-3)Man |
| α1-3, α1-3, α1-6-Mannopentaose | Manα1-6(Manα1-3)Manα1-6(Manα1-3)Man |
| **Fucosylated structures** | |
| Lacto-*N*-fucopentose I | Fucα1-2Galβ1-3Glc*N*Acβ1-3Galβ1-4Glc |
| Lacto-*N*-fucopentose II | Galβ1-3(Fuca1-4)Glc*N*Acβ1-3Galβ1-4Glc |
| Lacto-*N*-fucopentose III | Galβ1-4(Fuca1-3)Glc*N*Acβ1-3Galβ1-4Glc |
| Lacto-*N*-difucohexose I | Fucα1-2Galβ1-3(Fucα1-4)Glc*N*Acβ1-3Galβ1-4Glc |
| Lacto-*N*-difucohexose II | Galβ1-3(Fucα1-4)Glc*N*Acβ1-3Galβ1-4(Fucα1-3)Glc |
| H-disaccharide | Fucα1-2Gal |
| 2'-Fucosyllactose | Fucα1-2Galβ1-4Glc |
| 3'-Fucosyllactose | Galβ1-4(Fucα1-3)Glc |
| Lewis$^x$ | Galβ1-4(Fucα1-3)Glc*N*Ac |
| Lewis$^a$ | Galβ1-3(Fucα1-4)Glc*N*Ac |

**Cont. Supplemental Table 1**

| | |
|---|---|
| Blood Group A-trisaccharide | Gal*N*Acα1-3(Fucα1-2)Gal |
| Lactodifucotetrose | Fucα1-2Galβ1-4(Fucα1-3)Glc |
| Blood Group B-Trisaccharide | Galβ1-3(Fucα1-2)Gal |
| Lewis$^{y}$ | Fucα1-2Galβ1-4(Fucα1-3)Glc*N*Ac |
| Blood Group H Type II Trisaccharide | Fucα1-2Galβ1-3Glc*N*Ac |
| Lewisb tetrasaccharide | Fucα1-2Galβ1-3(Fucα1-4)Glc*N*Ac |
| Sulpho Lewis$^{a}$ | $SO_3$-3Galβ1-3(Fucα1-4)Glc*N*Ac |
| Sulpho Lewis$^{x}$ | $SO_3$-3Galβ1-4(Fucα1-3)Glc*N*Ac |
| Monofucosyl-para-Lacto-*N*-hexose IV | Galβ1-3Glc*N*Acβ1-3Galβ1-4(Fucα1-3)Glc*N*Acβ1-3Galβ1-4Glc |
| Monofucosyllacto-*N*-hexose III | Galβ1-4(Fucα1-3)Glc*N*Acβ1-6(Galβ1-3Glc*N*Acβ1-3)Galβ1-4Glc |
| Difucosyllacto-*N*-hexose | Galβ1-4(Fucα1-3)Glc*N*Acβ1-6(Fucα1-2Galβ1-3Glc*N*Acβ1-3)Galβ1-4Glc |
| Trifucosyllacto-*N*-hexose | Galβ1-4(Fucα1-3)Glc*N*Acβ1-6(Fucα1-2Galβ1-3(Fucα1-4)Glc*N*Acβ1-3)Galβ1-4Glc |
| Lacto-*N*-fucopentaose VI | Galβ1-4GlcNAcβ1-3Galβ1-4(Fucα1-3)Glc |
| Lacto-*N*-neodifucohexaose I | Fucα1-2Galβ1-4(Fucα1-3)GlcNAcβ1-3Galβ1-4Glc |
| Lacto-*N*-neodifucohexaose II | Fucα1-3Galβ1-4GlcNAcb1-3Galβ1-4(Fucα1-3)Glc |
| Trifucosyllacto-*N*-neoteraose I | Fucα1-2Galβ1-4(Fucα1-3)GlcNAcβ1-3(Fucα1-2)Galβ1-4Glc |
| Monofucosyllacto-*N*-neohexaose I | Galβ1-4(Fucα1-3)GlcNAcβ1-6(Galβ1-4GlcNAcβ1-3)Galβ1-4Glc |
| Difucosyllacto-*N*-neohexaose I | Galβ1-4(Fucα1-3)GlcNAcβ1-6(Galβ1-4(Fucα1-3)GlcNAcβ1-3)Galβ1-4Glc |
| Difucosyllacto-*N*-neohexaose II | Fucα1-2Galβ1-4(Fucα1-3)GlcNAcβ1-6(Galβ1-4GlcNAcβ1-3)Galβ1-4Glc |
| Monofucosyl(1-3)-iso-lacto-*N*-octaose | Galβ1-3GlcNAcβ1-3Galβ1-4(Fucα1-3)GlcNAcβ1-6(Galβ1-3GlcNAcβ1-3)Galβ1-4Glc |
| Trifucosyl(1-2,1-2,1-3)-iso-lacto-N-octaose | Fucα1-2Galβ1-3GlcNAcβ1-3Galβ1-4(Fucα1-3)GlcNAcβ1-6(Galβ1-3GlcNAcβ1-3)Galβ1-4Glc |
| Blood Group A Tetrasaccharide | GalNAcβ1-3(Fucα1-2)Galβ1-4Glc |
| Blood Group B pentasaccharide | Galβ1-3(Fucα1-2)Galβ1-4(Fucα1-3)Glc |
| **Neu5Ac containing structures** | |
| Sialyl Lewis$^{a}$ | Neu5Acα2-3Galβ1-3(Fucα1-4)Glc*N*Ac |
| Sialyl Lewis$^{x}$ | Neu5Acα2-3Galβ1-4(Fucα1-3)Glc*N*Ac |
| Sialyllacto-*N*-tetrose a | Neu5Acα2-3Galβ1-3Glc*N*Acβ1-3Galβ1-4Glc |
| Monosialyl, monofucosyllacto-*N*-neohexose | Galβ1-4(Fucα1-3)Glc*N*Acβ1-6(Neu5Acα2-6Galβ1-4Glc*N*Acβ1-3)Galβ1-4Glc |
| 2,3'-Sialyllactosamine | Neu5Acα2-3Galβ1-4Glc*N*Ac |
| 2,6'-Sialyllactosamine | Neu5Acα2-6Galβ1-4Glc*N*Ac |
| LS-Tetrasaccharide a | Neu5Acα2-3Galβ1-3GlcNAcβ1-3Galβ1-4Glc |

**Cont. Supplemental Table 1**

| | |
|---|---|
| LS-Tetrasaccharide b | Galβ1-3(Neu5Acα2-6)Glc*N*Acβ1-3Galβ1-4Glc |
| LS-Tetrasaccharide c | Neu5Acα2-6Galβ1-4Glc*N*Acβ1-3Galβ1-4Glc |
| Disialyllacto-*N*-tetrose | Neu5Acα2-3Galβ1-3(Neu5Acα2-6)Glc*N*Acβ1-3Galβ1-4Glc |
| 2,3'-Sialyllactose | Neu5Acα2-3Galβ1-4Glc |
| 2,6'-Sialyllactose | Neu5Acα2-6Galβ1-4Glc |
| Colominic acid | (Neu5Acα2-8Neu5Ac)n (n<50) |
| Biantennary 2,6-sialylated-*N*-glycan-Asn | Neu5Acα2-6Galβ1-4Glc*N*Acβ1-2Manα1-6(Neu5Acα2-6Galβ1-4Glc*N*Acβ1-2Manα1-6)Manβ1-4Glc*N*Acβ1-4Glc*N*Ac-Asn |
| **Carageenan and Glycoaminoglycans (GAGS)** | |
| Neocarratetrose-41, 3-di-*O*-sulphate ($Na^+$) | $C_{24}H_{36}O_{25}S_2Na_2$ (Mixed anomers. Tetrasaccharide of regular κ - carrageenan) |
| Neocarratetrose-41-*O*-sulphate ($Na^+$) | $C_{24}H_{37}O_{22}SNa$ (Mixed anomers. Derived from C1003 by removal of the non-reducing terminal 4-sulphate) |
| Neocarrahexose-24,41, 3, 5-tetra-*O*-sulphate ($Na^+$) | $C_{36}H_{52}O_{40}S_4Na_4$ (Mixed anomers. A hybrid sequence comprising carrageenan disaccharides in the order k-i-k, derived from the carrageenan from *Chondrus crispus*) |
| Neocarrahexose-41, 3, 5-tri-*O*-sulphate ($Na^+$) | $C_{36}H_{53}O_{37}S_3Na_3$ (Mixed anomers. Hexasaccharide of regular κ-carrageenan) |
| Neocarraoctose-41, 3, 5, 7-tetra-*O*-sulphate ($Na^+$) | $C_{48}H_{70}O_{49}S_4Na_4$ (Mixed anomers. Octasaccharide of regular κ-carrageenan) |
| Neocarradecose-41, 3, 5, 7, 9-penta-*O*-sulphate ($Na^+$) | $C_{60}H_{87}O_{61}S_5Na_5$ (Mixed anomers. Decasaccharide of regular κ- carrageenan) |
| ΔUA-2S → Glc*N*S-6S $Na_4$ (I-S) | $C_{12}H_{15}NO_{19}S_3Na_4$ (Predominant disaccharide produced from heparin by heparinase I and II) |
| ΔUA → Gluc*N*S-6S $Na_3$ (II-S) | $C_{12}H_{16}NO_{16}S_2Na_3$ (Produced from heparinase II digestion of heparin and heparin sulphate) |
| ΔUA → 2S-Glc*N*S $Na_3$ (III-S) | $C_{12}H_{16}NO_{16}S_2Na_3$ (Produced from heparin by digestion with heparinase I and II) |
| ΔUA → 2S-Glc*N*Ac-6S $Na_3$ (I-A) | $C_{14}H_{18}NO_{17}S_2Na_3$ (Minor component produced from heparin by heparinase II) |
| ΔUA → Glc*N*Ac-6S $Na_2$ (II-A) | $C_{14}H_{19}NO_{14}SNa_2$ (Product of the action of heparinases II and III on heparin and heparan sulphate) |
| ΔUA → 2S-Glc*N*Ac $Na_2$ (III-A) | $C_{14}H_{19}NO_{14}SNa_2$ (Minor product of the action of heparinase II on heparin) |
| ΔUA → Glc*N*Ac Na (IV-A) | $C_{14}H_{20}NO_{11}Na$ (Produced from heparin sulphate by digestion With heparinase III) |
| ΔUA → Gal*N*Ac-4S $Na_2$ (ΔDi-4S) | $C_{14}H_{19}NO_{14}SNa_2$ (Produced from various chondroitin sulphates By the action of chondroitinases ABC, B and AC-1) |

**Cont. Supplemental Table 1**

| | |
|---|---|
| ΔUA → Gal*N*Ac-6S Na2 (ΔDi-6S) | $C_{14}H_{19}NO_{14}SNa_2$ (Produced from various chondroitin sulphates By the action of chondroitinases ABC, AC-1 and C) |
| ΔUA → Gal*N*Ac-4S,6S $Na_3$ (ΔDi-disE) | $C_{14}H_{18}NO_{17}S_2Na_3$ (Produced from various chondroitin sulphates By the action of chondroitinases ABC, B and AC-1) |
| ΔUA → 2S-Gal*N*Ac-4S $Na_2$ (ΔDi-disB) | $C_{14}H_{18}NO_{17}S_2Na_3$ (Produced from various chondroitin sulphates by action of chondroitinase ABC and/or B. Most typically from chondroitin sulphate B (dermatan sulphate)) |
| ΔUA → 2S-Gal*N*Ac-6S $Na_3$ (ΔDi-disD) | $C_{14}H_{18}NO_{17}S_2Na_3$ (Produced from various chondroitin sulphates by the action of chondroitinase ABC) |
| ΔUA → 2S-Gal*N*Ac-4S-6S $Na_4$ (ΔDi-tisS) | $C_{14}H_{17}NO_{20}S_3Na_4$ (Produced as a minor component by the action of chondroitinase ABC on various chondroitin sulphates, particularly B) |
| ΔUA → 2S-Gal*N*Ac-6S $Na_2$ (ΔDi-UA2S) | $C_{14}H_{19}NO_{14}SNa_2$ (Produced as a minor component from various chondroitin sulphates by the action of chondroitinase ABC) |
| ΔUA → Glc*N*Ac Na (ΔDi-HA) | $C_{14}H_{20}NO_{11}Na$ (The only unsaturated disaccharide produced from hyaluronic acid by the action of chondroitinase ABC or AC-1) |
| Hyaluronan fragments (4mer) | (GlcAβ1-3Glc*N*Acβ1-4)n (n=4) |
| Hyaluronan fragments (8mer) | (GlcAβ1-3Glc*N*Acβ1-4)n (n=8) |
| Hyaluronan fragments (10mer) | (GlcAβ1-3Glc*N*Acβ1-4)n (n=10) |
| Hyaluronan fragments (12mer) | (GlcAβ1-3Glc*N*Acβ1-4)n (n=12) |
| Heparin | (GlcA/IdoAα/β1-4Glc*N*Acα1-4)n (n=200) |
| Chondroitin sulfate | (GlcA/IdoAβ1-3(±4/6S)Gal*N*Acβ1-4)n (n<250) |
| Dermatan sulfate | ((±2S)GlcA/IdoAα/b1-3(±4S)Gal*N*Acβ1-4)n (n<250) |
| Chondroitin 6-Sulfate | (GlcA/IdoAβ1-3(±6S)Gal*N*Acβ1-4)n (n<250) |
| HA - 4 | (GlcAβ1-3Glc*N*Acβ1-4)n (n=4) |
| HA - 6 | (GlcAβ1-3Glc*N*Acβ1-4)n (n=6) |
| HA - 8 | (GlcAβ1-3Glc*N*Acβ1-4)n (n=8) |
| HA 10 | (GlcAβ1-3Glc*N*Acβ1-4)n (n=10) |
| HA-12 | (GlcAβ1-3Glc*N*Acβ1-4)n (n=12) |
| HA-14 | (GlcAβ1-3Glc*N*Acβ1-4)n (n=14) |
| HA-16 | (GlcAβ1-3Glc*N*Acβ1-4)n (n=16) |
| HA 30000 Da | (GlcAβ1-3Glc*N*Acβ1-4)n |
| HA 107000 Da | (GlcAβ1-3Glc*N*Acβ1-4)n |

**Supplemental Table 2- Clusters A-H and their interacting amino acids.**

| Cluster | Amino acids |
|---|---|
| **A** | **PHE-203**, ILE-205, VAL-211, ILE-224, **TYR-228**, VAL-231, ALA-234, THR-235, VAL-238, LEU-250, TYR-252, LEU-263, VAL-265, ALA-285, **ILE-287** |
| B | GLY-206, VAL-207, LYS-208, LEU-242, GLU-243, LYS-269, LEU-280, ASN-281 |
| C | VAL-60, ASN-63, THR-64, SER-67, ALA-94, ASN-95, SER-96, HIS-97 |
| **D** | SER-102, MSE-103, PHE-104, THR-114, GLU-126, ASN-133, **ALA-136**, ARG-153, **TYR-151**, LEU-167, ALA-181, LEU-182, MET-183 |
| E | PHE-105, LYS-106, ASN-107, ARG-108, ASN-172, VAL-179 |
| F | LYS-138, ASN-142, GLU-144, ILE-145, SER-146, SER-276, LYS-277, ASP-278 |
| G | PRO-76, LYS-77, ASP-78, THR-79, LYS-84, PHE-105, ASN-107, ARG-108, LEU-111, ASN-172, GLU-173, VAL-179 |
| H | SER-96, HIS-97, VAL-98, ALA-99, ARG-117, ASP-118, MSE-155, PRO-156, ASN-157, ALA-159, VAL-161, SER-187, SER-190 |

**Supplemental Table 3- TlpALBD docking analysis with fumarate and arginine.**

| | **Fumarate** | **Arginine** |
|---|---|---|
| **Cluster** | Cluster Occupation % | Cluster Occupation % |
| **A (membrane-proximal Cache)** | **45** | 25 |
| B | 10 | 5 |
| C | 20 | 5 |
| **D (membrane-distal Cache)** | 10 | **55** |
| E | 5 | - |
| F | 10 | - |
| G | - | 5 |
| H | - | 5 |

**Supplemental Table 4- Primers for site directed mutagenesis of TlpA$_{LBD}$.** Mutated codons, bolded; restriction sites used for screening clones, underlined.

| **Primer name** | **Sequence (5' → 3')** |
|---|---|
| TlpA_D165A_For | GGGGCGGAAGTTTATGGAGTT**GCT**ATTCTTTTACCTTTATTG |
| TlpA_D165A_Rev | CAATAAAGGTAAAAGAAT**AGC**AACTCCATAAACTTCCGCCCC |
| TlpA_M183A_For | GAGGTTGTAGGGGCTTTG**GCG**GTTTTTATTTCCATTGACAGC |
| TlpA_M183A_Rev | GCTGTCAATGGAAATAAAAAC**CGC**CAAAGCCCCTACAACCTC |
| TlpA_Y228A_For | GACAAACCTATCGCAGAAATT**GCT**AAGAGCGTACCTAAAGCC |
| TlpA_Y228A_Rev | GGCTTTAGGTACGCTCTT**AGC**AATTTCTGCGATAGGTTTGTC |
| TlpA_Y252A_For | CTCTAAAGCGACTTTAG<u>AA**GCT**T</u>TAGATCCCTTTAGCCATAAGG |
| TlpA_Y252A_Rev | CCTTATGGCTAAAGGGATCTA<u>A**AGC**TT</u>CTAAAGTCGCTTTAGAG |
| TlpA_D254A_For | CTAAAGC<u>GACTC</u>TAGAATACTTA**GCT**CCCTTTAGCCATAAGG |
| TlpA_D254A_Rev | CCTTATGGCTAAAGGG**AGC**TAAGTATTCTA<u>GAGTC</u>GCTTTAG |
